# Transcriptional coupling of metabolism and cell mechanics controls human neuroepithelium morphogenesis

**DOI:** 10.64898/2026.09.22.753382

**Authors:** Camil Mirdass, Lucas Denis, Hugo Lachuer, Mikaëlle Bocel, Anis Bourou, Kamal Bouhali, Nicolas Valentin, Sandrine Adiba, Alice Marteil, Céline Chollet, Emeline Chu-Van, Xavier Baudin, Catherine Pioche-Durieu, Sarah Chebouti, Bruno Estebe, Frédéric Relaix, Florence Castelli, Valérie Dupe, Sébastien Léon, Stéphane Nedelec, Nicolas Borghi, Vanessa Ribes

## Abstract

Tissue morphogenesis integrates developmental programs with cell mechanics, yet the mechanisms coupling these processes remain poorly understood. We investigated this link during neuroepithelial morphogenesis in human iPSC-derived dorsal spinal organoids. We found that PAX3, a transcription factor whose loss causes neural tube defects, promotes force-bearing adherens junction assembly. It does so by driving apical clustering of N-cadherin complexes, thereby enabling neuroepithelial morphogenesis, a function conserved in the mouse neural tube. Transcriptomic, metabolomic, and functional analyses show that PAX3 sustains upper glycolysis and the associated mannose-dependent glycosylation pathways. Exogenous mannose rescues cadherin clustering and adherens junction mechanics following *PAX3* loss, whereas inhibition of glycolysis or glycosylation recapitulates these defects. These findings establish mannose metabolism as a PAX3-regulated pathway conferring mechanical competence to developing tissues.

## Introduction

During embryogenesis, tissue morphogenesis emerges, in part, from the conversion of developmental programs into cell mechanical properties that shape tissues (*1*, *2*). Yet, the molecular mechanisms underlying this process have remained particularly difficult to define in amniotes. Measuring tissue mechanics *in vivo* remains technically challenging because morphogenesis unfolds over prolonged developmental timescales within tissues embedded deep inside the embryo (*3*). Moreover, the pleiotropic nature of the transcription factors that orchestrate developmental programs has made it difficult to distinguish the molecular cascades directly controlling tissue morphogenesis from those underlying other developmental processes (*4*).

Vertebrate neurulation provides a compelling paradigm in which to address this question (*5*, *6*). During this morphogenetic process, an elongated tubular pseudostratified neuroepithelium is assembled and remodeled to form the spinal cord primordium (*5*). In amniotes, the spinal neural tube forms through two distinct morphogenetic mechanisms: neural plate folding anteriorly during primary neurulation and cavitation of the neuralized tail bud posteriorly during secondary neurulation (*5*). Failure of either process results in spinal dysraphisms (spina bifida)(*5*, *6*). Genetic studies in humans and animal models have implicated hundreds of genes in these morphogenetic processes (*5–9*). Although disruption of these genes invariably results in spinal dysraphisms, they span remarkably diverse functional classes, including transcription factors, chromatin modifiers, signaling molecules, metabolic enzymes, and regulators of cell adhesion and cytoskeletal organization.

Transcription factors associated with spinal dysraphisms provide a unique genetic entry point to identify such mechanisms (*5*). Among them, we focused on PAX3, whose requirement for spinal neurulation is conserved in humans and mice (*5*). In amniotes, PAX3 is expressed in progenitors of the future dorsal spinal neuroepithelium throughout neurulation (*10–13*). In mice, complete loss of *Pax3* causes fully penetrant failure of caudal neural tube closure, whereas heterozygous embryos develop similar defects at lower, incomplete penetrance (∼10%), demonstrating a strong dosage dependence (*14–16*). In humans, heterozygous loss-of-function variants are primarily associated with Waardenburg syndrome (*17*) and, more rarely, with open spinal dysraphisms, up to the most severe form, myelomeningocele (*7*, *9*, *18–22*).

Despite compelling genetic evidence implicating PAX3 in spinal neurulation, the gene regulatory network through which it controls neural tube closure remains unknown. Previous studies in animal models have primarily focused on its roles in dorsoventral patterning and neural crest specification, where Pax3 acts as both an activator and a repressor in a spatiotemporally regulated manner (*23–28*). Genetic complementation experiments demonstrated that replacing one *Pax3*-null allele with a constitutively active *PAX3-FOXO1* allele was sufficient to rescue spinal neurulation, indicating that this morphogenetic function primarily depends on Pax3-mediated transcriptional activation (*29*). An important clue comes from the long-recognized folate responsiveness of *Pax3*-deficient embryos, one of the few mouse models in which spinal dysraphisms are partially rescued by maternal folic acid supplementation and exacerbated by folate deficiency (*15*, *30–33*). *Pax3* deficiency does not overtly alter folate metabolism. Instead, folic acid has been proposed to rescue neurulation by bypassing the defect in *de novo* purine and pyrimidine biosynthesis caused by *Pax3* loss (*15*, *30*, *32*). Together, these observations suggested that PAX3 regulates a metabolic program required for spinal neurulation, although the underlying mechanism remained unresolved.

## Results

### Stereotypical neuroepithelial morphogenesis in human dorsal spinal organoids

To investigate how PAX3 regulates neuroepithelial morphogenesis, we used a previously established human iPSC-derived dorsal spinal organoid model that recapitulates key features of embryonic dorsal spinal cord development and enables quantitative analysis of neuroepithelium-intrinsic morphogenesis (*34*). Organoids were patterned toward dorsal spinal identity by temporally controlled Wnt (via the GSK3 inhibitor CHIR99021), retinoic acid (RA), and BMP (BMP4) signaling (Fig. 1A). ROCK inhibition (Y27632) was applied during the first two days to promote survival. At day 7, organoids were composed predominantly of SOX1⁺ neural progenitors (∼90%), with ∼75% expressing PAX6 (Fig. 1B and fig. S1B). Dorsal identity was robustly established, as indicated by widespread PAX3 and PAX7 expression (∼80% of cells; Fig. 1, B to D). Approximately 40% of progenitors were OLIG3⁺, consistent with specification toward the dorsal-most interneuron progenitor domains (dP1-dP3), while a minor population (<5%) of SOX10⁺ neural crest cells (NCCs) localized to the organoid periphery (Fig. 1B and fig. S1B).

**Fig. 1.**
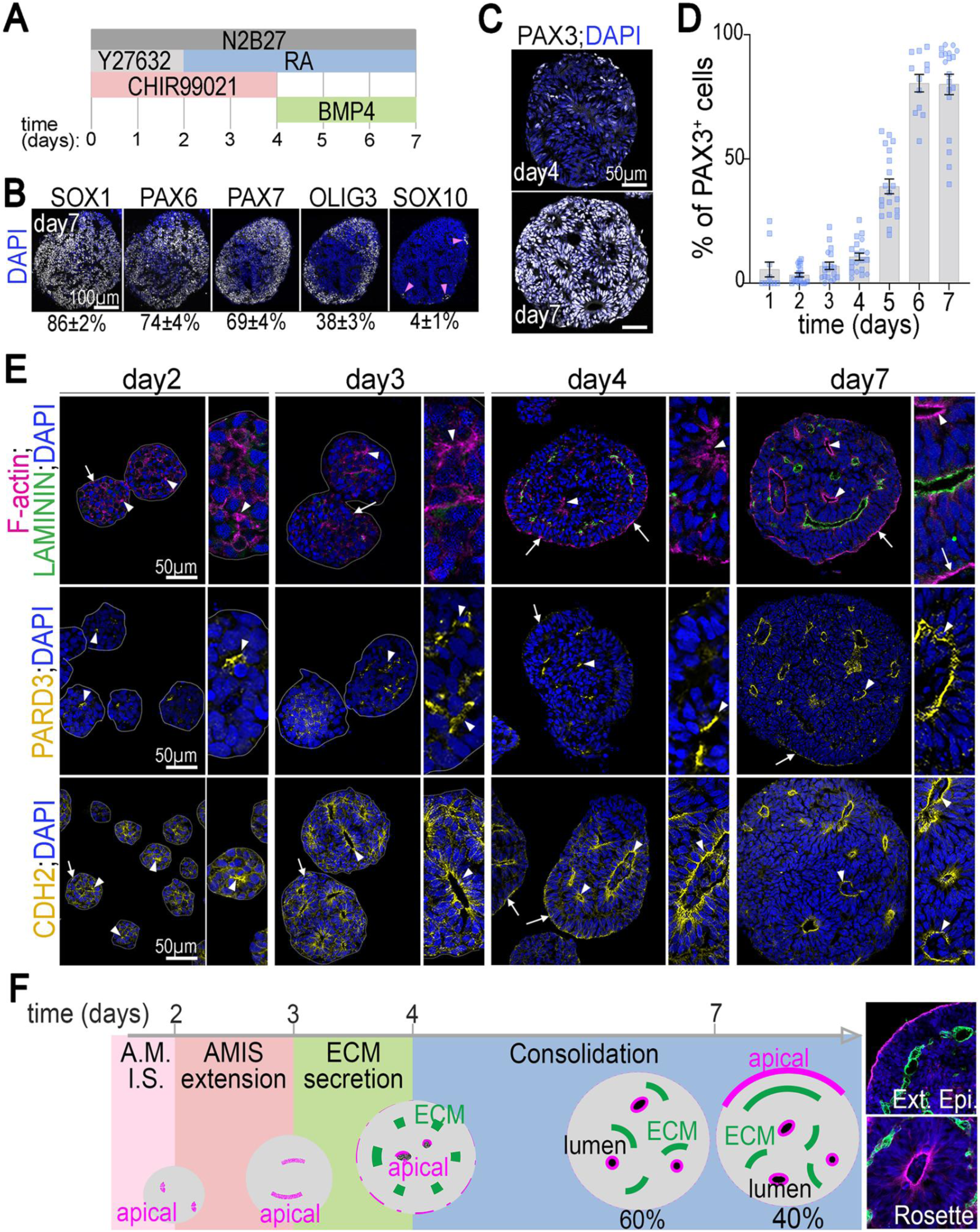
Spatiotemporal emergence of epithelial rosettes and peripheral epithelia in dorsal PAX3^+^ spinal cord organoids. **(A)** Schematic of the differentiation of human iPSCs into dorsal spinal cord organoids. **(B and C)** Representative images of day 7 wild-type (WT) organoids stained for markers of dorsal spinal cord identity and for DAPI. Mean percentage of marker-positive cells ± SEM is given below each image in (B). Pink arrowheads, SOX10^+^ neural crest cells. **(D)** Percentage of PAX3^+^ cells in WT organoids at the indicated differentiation time points. Dots, individual organoids (one section per organoid, symbol shape indicates WT iPSC line); bars, mean ± SEM. **(E)** Representative images of day 2, 3, 4, and 7 WT organoids stained for LAMININ, PARD3, CDH2 (N-cadherin), F-actin (phalloidin), and DAPI. Arrowheads, cell-cell adhesions of nascent or established inner rosettes; arrows, adhesions of peripheral epithelia. **(F)** Proposed model for the stepwise formation of inner rosettes and peripheral epithelia. Percentages indicate organoids containing inner rosettes only (60%) or both inner rosettes and peripheral epithelia (40%). Higher-magnification images stained for LAMININ, F-actin (phalloidin), and DAPI are shown at right. Color code: pink, apical surfaces; green, extracellular matrix (ECM); black, lumen. Sample sizes and statistical analyses are provided in Data S1.

PAX3 expression emerged at day 4 and became widespread by day 7 (Fig. 1, C and D). During this developmental window, progenitors progressively self-organized into multiple polarized pseudostratified neuroepithelial compartments, each delimited by LAMININ-rich extracellular matrix (ECM) and displaying apical enrichment of F-actin cytoskeleton, the polarity protein PARD3 and the adhesion protein N-cadherin (CDH2), recapitulating the characteristic apico-basal organization of the neural tube neuroepithelium (*35*) (Fig. 1, E and F). Time-course analysis showed that, as early as day 2 and prior to detectable ECM deposition, cells established F-actin⁺/PARD3⁺/CDH2⁺ cell-cell contacts reminiscent of apical membrane initiation sites (AMIS) (*36*, *37*), both internally and at the organoid surface (Fig. 1, E and F). By day 3, tissue organization did not change, although these apical domains had increased in size (Fig. 1, E and F). From day 4 onwards, cell organization changed markedly. Progressive LAMININ deposition delineated emerging epithelial compartments, initially appearing as discrete puncta before assembling into elongated LAMININ-positive stretches along the basal surface of developing epithelia (Fig. 1, E and F). Concomitantly, apico-basal polarity strengthened progressively, with increasing apical enrichment of CDH2, PARD3 and F-actin (Fig. 1, E and F). During this period, two distinct epithelial architectures emerged. Cells within the organoid interior organized into neuroepithelial rosettes surrounding central lumens, present in all organoids by day 7 (arrowheads in Fig. 1, E and F). In a subset of organoids (∼40%), cells located at the organoid surface additionally formed peripheral epithelia whose apical surface faced the external environment (arrows in Fig. 1, E and F). Together, these observations establish a reproducible model of neuroepithelial morphogenesis that recapitulates cavitation during neural organoid development and secondary neurulation *in vivo* (*38*).

### PAX3 dosage-dependent neuroepithelial organization in human spinal organoids

We generated an isogenic allelic series of human iPSCs carrying heterozygous or homozygous CRISPR-Cas9-induced frameshift null mutations in *PAX3* (3 to 4 independent clones per genotype, see Materials and Methods section) (fig. S2A). Mutant organoids exhibited the expected gene-dosage-dependent reduction or complete loss of PAX3 protein while retaining competence to generate dorsal spinal progenitors (Fig. 2, A and B, and fig. S2, B and C). To test whether our organoid system reproduces established functions of PAX3 (*15*, *23–27*), we first examined neural specification and progenitor proliferation in day 7 organoids. *PAX3* loss did not alter the overall rostrocaudal identity of spinal organoids, which maintained a cervical-to-anterior brachial identity (*HOXA5*⁺; *HOXC5*⁺; *HOXC6*⁺; *HOXC8*⁻) across genotypes (fig. S1A and S2D). Likewise, dorsal spinal specification was largely preserved, with sustained expression of PAX7 and OLIG3 (Fig. 2, A and B and fig. S2 C). The most prominent specification defects were loss of SOX10⁺ neural crest cells and increased *DBX1* expression, indicative of mild ventralization (Fig. 2B and fig. S2, C and D). *PAX3*^−/−^ organoids also exhibited reduced progenitor expansion, reflected by smaller organoid size and reduced G2/M-phase fractions (Fig. 2B and fig. S2, E and F). Heterozygous organoids remained largely comparable to WT controls, although several parameters, including neural crest cell abundance and proliferative indices, showed intermediate values (Fig. 2B and fig. S2, D to F). Overall, these findings indicate that PAX3 is largely dispensable for dorsal spinal specification during this stage of development (*15*, *23*, *24*), enabling analysis of its morphogenetic functions independently of major changes in neural identity.

**Fig. 2.**
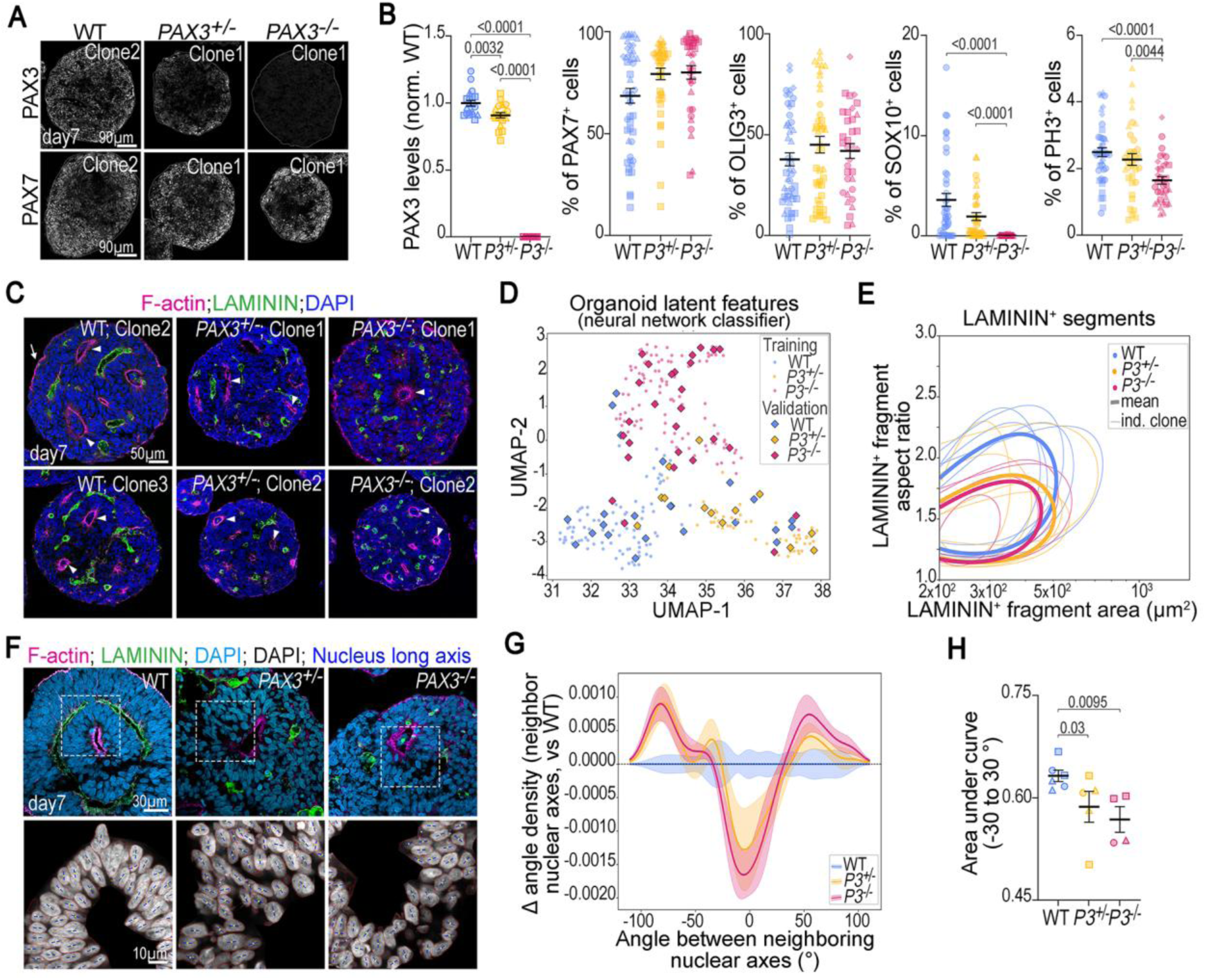
Neuroepithelial architecture defects revealed by neural network-based and quantitative analyses following progressive loss of *PAX3* alleles. (A) Representative images of day 7 WT, *PAX3*^+/−^, and *PAX3*^−/−^ organoids stained for PAX7, PAX3, and DAPI. **(B)** Nuclear PAX3 fluorescence intensity and percentages of PAX7^+^, OLIG3^+^, SOX10^+^, and phospho-histone H3 (PH3)^+^ nuclei in day 7 WT, *PAX3*^+/−^, and *PAX3*^−/−^ organoids. Dots, individual organoids (symbol shape indicates clone); bars, mean ± SEM across independent organoids. **(C)** Representative images of day 7 WT and *PAX3*^−/−^ organoids (two clones per genotype) stained for LAMININ, F-actin (phalloidin), and DAPI. Arrowheads, cell-cell adhesions of inner rosettes; arrows, adhesions of peripheral epithelia. **(D)** UMAP of latent features from a ResNet-50 classifier trained on images of day 7 WT, *PAX3*^+/−^, and *PAX3*^−/−^ organoids stained for LAMININ, F-actin, and DAPI. Each point, one organoid; circles, training dataset; diamonds, independent hold-out validation dataset. **(E)** Contours at 70% of the maximum kernel density estimate for LAMININ fragment aspect ratio versus fragment area (from fig. S3E). Thick contours, genotype means; thin contours, individual clonal replicates. **(F)** Representative images of day 7 WT and *PAX3*^−/−^ organoids stained for LAMININ, F-actin (phalloidin), and DAPI (top); dashed boxes, regions shown at higher magnification (bottom). Bottom, DAPI channel alone, with segmented nuclei outlined in red and nuclear long axes in blue. **(G)** Difference from WT in the population distribution of nearest-neighbor nuclear alignment angles: for each angular bin, the proportion of nuclei of the indicated genotype minus that of WT. Values, mean ± SEM. **(H)** Probability of nearest-neighbor nuclear alignment angles between −30° and +30°. Dots, independent experiments (symbol shape indicates clone); bars, mean ± SEM across independent experiments. Mann-Whitney P values are shown on the plots; individual data points, sample sizes, and statistical analyses are provided in Data S1.

We next investigated whether neuroepithelial organization was sensitive to PAX3 dosage by analyzing LAMININ, F-actin and DAPI staining at day 7, which together report epithelial architecture and compartmentalization within organoids (Fig. 2C and fig. S3A). Across independent experiments and clones, *PAX3*^−/−^ organoids were readily distinguishable from WT controls, displaying reduced epithelial compartmentalization, loss of peripheral epithelial structures and disorganized neuroepithelial rosettes (Fig. 1E and 2C and fig. S3, A and B). In contrast, *PAX3*^+/−^ organoids displayed greater morphological heterogeneity, with morphologies ranging from WT-like to *PAX3*^−/−^-like, precluding the definition of a stereotypical heterozygous phenotype at the single organoid level. However, scoring of peripheral epithelial structures at the organoid population level positioned heterozygous organoids in an intermediate position between WT and *PAX3*^−/−^ organoids (fig. S3B).

To further assess the penetrance of these architectural abnormalities in a feature-agnostic manner, we applied deep-learning-based image classification to 310 histological sections, each derived from an individual organoid generated across three to four independent clones and eight differentiation experiments per genotype. Sections stained for LAMININ, F-actin and nuclei comprised 114 WT, 65 *PAX3*^+/−^ and 131 *PAX3*^−/−^ organoids. Images were randomly partitioned into training (80%) and independent validation (20%) datasets. A ResNet-50 classifier showed progressive reductions in prediction error in both datasets, indicating effective learning with limited overfitting (fig. S3C). The classifier correctly identified the majority of previously unseen images, achieving the highest accuracy for *PAX3*^−/−^ organoids (85.2%), while classification performance was lower for WT (56.5%) and *PAX3*^+/−^ organoids (61.5%) (fig. S3D). UMAP projection of latent image features revealed clear genotype-associated segregation, with validation images intermingled with training samples, indicating that the learned morphological signatures generalised across independent datasets (Fig. 2D). We therefore asked whether this genotype-associated segregation was underpinned by measurable morphological traits, focusing on two features: (i) extracellular matrix (ECM) geometry and (ii) cellular organization within neuroepithelial rosettes. Analysis of LAMININ-positive fragment area and aspect ratio distributions, visualized by kernel density estimation, revealed a progressive shift toward smaller and less elongated structures with decreasing PAX3 dosage (Fig. 2E and fig. S3E). Alignment angles between neighboring nuclei within neuroepithelial rosettes as a proxy for local epithelial order, were shifted in *PAX3*^−/−^ organoids, which showed fewer low-angle pairs and more large-angle pairs than WT controls (Fig. 2, F to H). *PAX3*^+/−^ organoids displayed intermediate phenotypes in both analyses, although with substantial inter-clone and experimental variability (Fig. 2, E, G and H).

Together, these findings establish dosage-sensitive defects in neuroepithelial morphogenesis in *PAX3*-mutant human spinal organoids despite largely preserved neural specification. This system provides an opportunity to dissect the molecular and mechanical mechanisms underlying PAX3-dependent neuroepithelial morphogenesis.

### *PAX3-dependent spatial restriction of adherens junction and actin assemblies to the apical* end of the lateral membrane

The epithelial disorganization, together with reduced cellular alignment within *PAX3*-deficient neuroepithelial rosettes, prompted us to examine the integrity of the apical junctional machinery involved in maintaining epithelial cohesion and architecture. To this end, we assessed apical junctional architecture in day 7 organoids by transmission electron microscopy (TEM) and high-resolution Airyscan imaging of junctional proteins (Fig. 3, A to D and fig. S4 and S5).

**Fig. 3.**
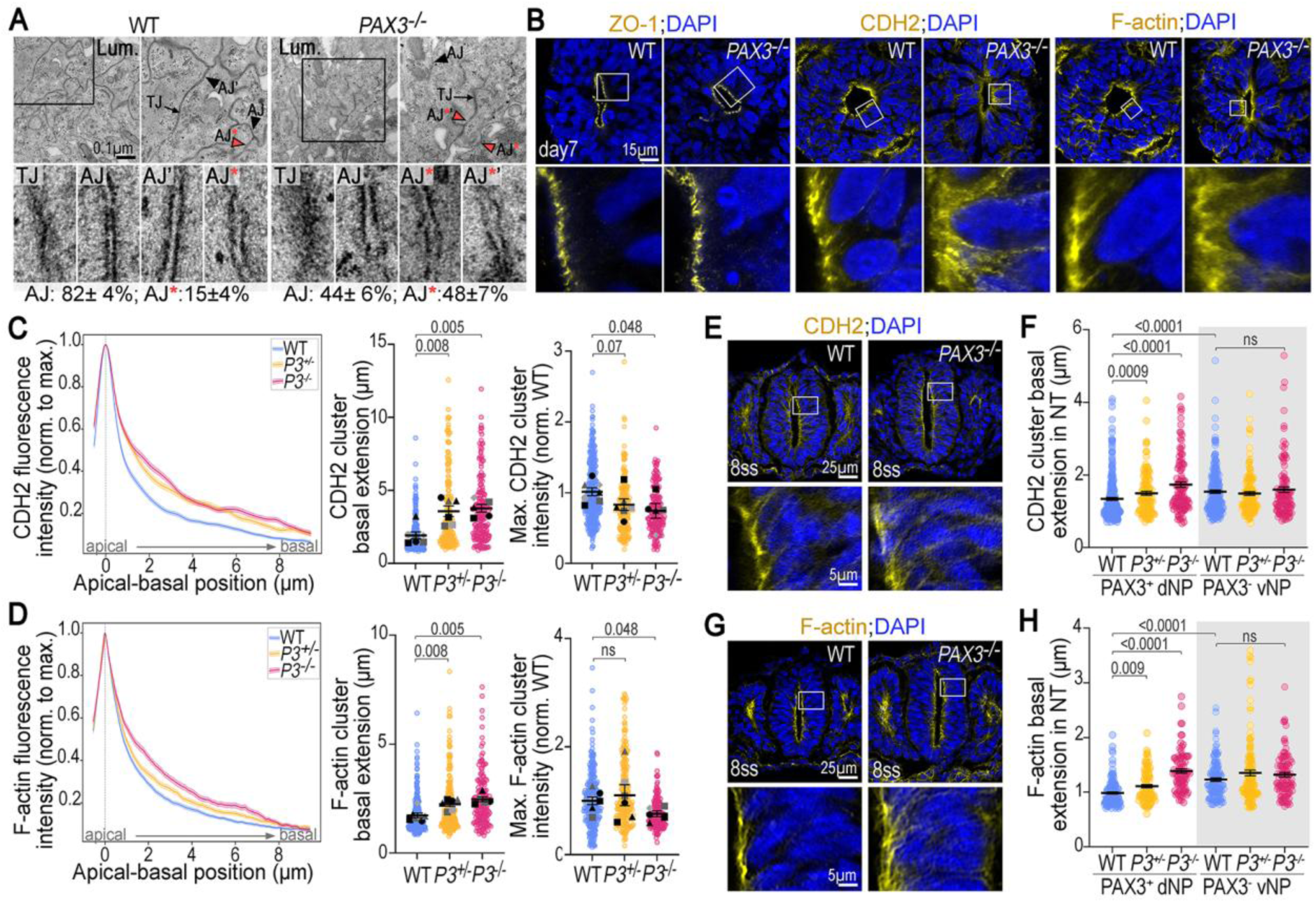
Altered apical adherens junction organization and N-cadherin clustering in *PAX3*-deficient human and mouse neural epithelia. (A to D) Day 7 human dorsal spinal cord organoids. **(E to H)** E8.5 mouse embryos (7 to 8 somites). **(A)** Representative TEM images centered on the apical surface of inner rosettes in WT and *PAX3*^−/−^ organoids. For each genotype: top left, low-magnification overview with the lumen indicated and a black box marking the region enlarged at top right; top right, the boxed region, in which one tight junction (TJ) and three adherens junctions (AJs) are highlighted; bottom, higher-magnification views of the highlighted TJ and AJs. Black arrows, TJs; arrowheads, AJs. Two AJ morphologies were identified: electron-dense, ladder-like junctions (AJ; black arrowheads) and less electron-dense junctions (AJ*; pink arrowheads outlined in black). Percentages indicate the proportion of AJs of each morphology per genotype. **(B)** Representative images of WT and *PAX3*^−/−^ organoid rosettes stained for ZO-1, CDH2 (N-cadherin), F-actin (phalloidin), and DAPI; white boxes, regions shown at higher magnification. **(C and D)** Apical CDH2 (C) and F-actin (D) signals within apical clusters in rosettes from day 7 WT, *PAX3*^+/−^, and *PAX3*^−/−^ organoids. Left to right: fluorescence intensity profiles along the apico-basal axis, normalized to the maximum; cluster basal extension; maximum cluster intensity normalized to WT. Colored dots, individual cell contacts; gray dots, per-experiment means (symbol shape indicates clone); bars, mean ± SEM across independent clone/experiment replicates. For intensity profiles: thick lines, mean across all measurements; thin lines, SEM across independent clones. **(E and G)** Representative images of WT and *Pax3^GFP/GFP^* embryos at somite level 5, stained for CDH2 (E) or F-actin (G), and DAPI; white boxes, regions shown at higher magnification. **(F and H)** Apical CDH2 (F) and F-actin (H) signals within apical clusters in embryos of the indicated genotypes, measured in the dorsal half (white background) or ventral half (gray background) of the neural tube. Dots, individual clusters (*n* = 4 embryos per genotype); bars, mean ± SEM across independent cross-sections. Mann-Whitney *P* values are shown on the plots; individual data points, sample sizes, and statistical analyses are provided in Data S1.

At the ultrastructural level, apical tight junctions (TJs) appeared largely preserved in *PAX3*-deficient rosettes (Fig. 3A). They were readily identified at the luminal apex, and the apical clustering of the TJ scaffold protein ZO-1 and PARD3 were unchanged (Fig. 3B and fig. S4, A and B). In contrast, TEM revealed marked alterations in adherens junction (AJ) morphology. We identified canonical AJs that displayed the characteristic electron-dense, ladder-like organization, and altered AJs (AJ*) that exhibited reduced electron density (Fig. 3A and fig. S5A). Quantitative classification showed a striking shift toward AJ* configurations in *PAX3*^−/−^ rosettes, whereas canonical AJs predominated in controls (fig. S5B). Consistent with these ultrastructural defects, Airyscan imaging revealed a profound reorganization of AJ components. In WT epithelia, CDH2 formed compact, high-intensity apical clusters extending on average 1.8 µm along the apico-basal axis (Fig. 3, B and C and fig. S4C). By contrast, *PAX3*-deficient rosettes exhibited broadened CDH2 domains that extended approximately twofold further along the apico-basal axis and displayed reduced peak fluorescence intensity (Fig. 3, B and C and fig. S4C). Comparable alterations were observed for α-catenin and β-catenin, indicating coordinated disorganization of the cadherin-catenin complex following *PAX3* loss (fig. S4, A and B).

Given the intimate coupling between AJs and the cortical actin cytoskeleton (*39*), we next examined apical F-actin organization (Fig. 3, B and D). In WT neuroepithelia, F-actin showed a compact apical enrichment extending on average 1.7 µm along the apico-basal axis (Fig. 3, B and D). In *PAX3*-deficient organoids, this apical actin enrichment remained detectable but was significantly broadened, extending to approximately 2.5 µm, and displayed a ∼1.3-fold reduction in peak fluorescence intensity (Fig. 3, B and D). Notably, *PAX3*^+/−^ organoids exhibited intermediate apical F-actin broadening relative to WT and *PAX3*^−/−^ organoids (Fig. 3, B and D). A similar dosage-dependent trend was observed for all adherens junction components analyzed, including CDH2, α-catenin and β-catenin (Fig. 3, B and C and fig. S4, A to C).

To determine whether the junctional defects identified in organoids were also present *in vivo*, we examined the neural tube of *Pax3*-mutant mouse embryos (Fig. 3, E to H and Fig. S4, D and E). Analyses were performed at the level of somite 5 in 7- to 8-somite embryos, where the neural tube was closed in controls and, in mutants, either closed but malformed or partially open. Analyses were carried out separately in the dorsal and ventral domains to account for regional variation in CDH2 and F-actin distribution and for the dorsal-restricted expression of Pax3 at this stage (Fig. 3, F and H, and fig. S4E). Consistent with the organoid phenotype, CDH2, β-catenin and F-actin exhibited approximately 1.3-fold broader apico-basal distributions in the dorsal neuroepithelium of *Pax3*-mutant embryos, whereas heterozygous embryos displayed intermediate phenotypes (Fig. 3, E to H and fig. S4, D and E). No significant alterations were detected in ventral neural tube regions lacking *Pax3* expression. These observations support a conserved physiological role for PAX3 in promoting the apical clustering and spatial confinement of junctional and actin assemblies during neural tube morphogenesis, conserved between human and mouse. Differences in epithelial topology between organoid rosettes and the embryonic neural tube may contribute to the stronger phenotype observed *in vitro*. The highly constricted apical surfaces of rosettes may be particularly sensitive to the out-of-plane broadening of junctional and actin assemblies that accompanies *PAX3* loss.

These observations identify PAX3 as a regulator of apical AJ and actin organization in spinal neuroepithelia, while overall apico-basal polarity remains largely intact. PAX3 confines cadherin-catenin and actomyosin assemblies to the apical extremity of the lateral domain; hence, loss of this confinement may explain the progressive disruption of epithelial architecture observed in *PAX3*-deficient organoids.

### PAX3 maintenance of actomyosin-dependent tension at adherens junctions

As AJs function as mechanosensitive hubs in epithelial tissues (*40*), we next investigated whether AJ mechanics were altered in *PAX3*-mutant neuroepithelia. We first examined the localisation of VINCULIN, an adaptor recruited to force-bearing AJs through actomyosin-dependent activation of α-catenin (*41*) (Fig. 4A and fig. S6A). In WT rosettes, VINCULIN formed discrete apical puncta with an average area of 0.16 µm² and occupied a narrower domain along the apico-basal axis than CDH2 (Fig. 4, A and B and fig. S6A). This distribution suggests that only a subset of cadherin-based adhesions is under sufficient mechanical load to recruit VINCULIN. In *PAX3*-deficient neuroepithelia, VINCULIN-positive clusters were markedly broadened, expanding to an average area of 0.45 µm² (Fig. 4, A and B and fig. S6A). Total VINCULIN signal per cluster remained largely unchanged (fig. S6B), whereas mean fluorescence intensity was reduced by approximately 50% (Fig. 4, A and B), indicating a redistribution of force-dependent VINCULIN recruitment across an expanded membrane domain.

**Fig. 4.**
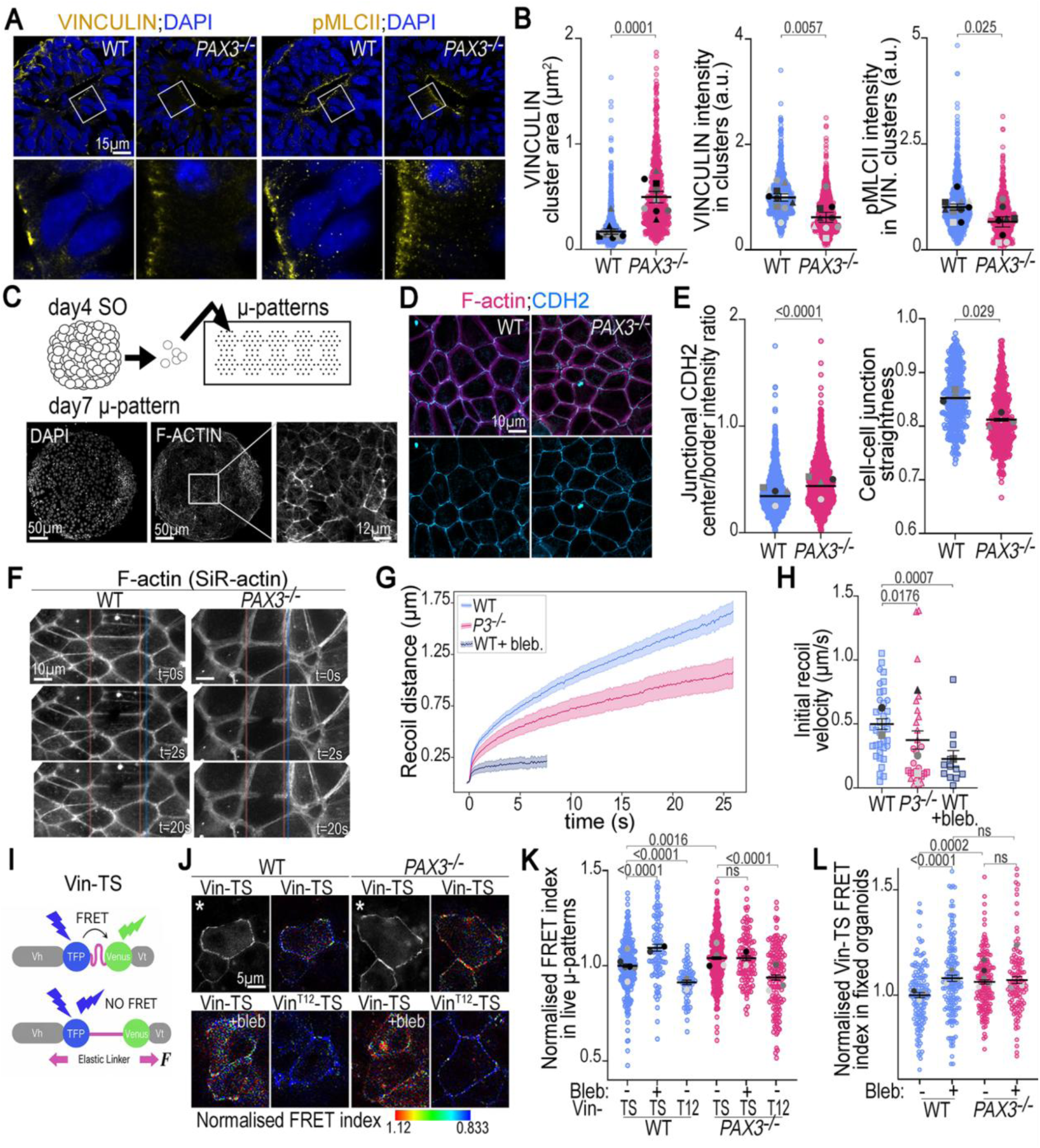
Impaired tension transmission and mechanotransduction at apical cell-cell junctions in *PAX3*-mutant spinal neural progenitors. (A) Representative images of day 7 WT and *PAX3*^−/−^ organoid rosettes stained for VINCULIN, phosphorylated myosin light chain II (pMLCII), and DAPI; white boxes, regions shown at higher magnification. **(B)** Apical VINCULIN and pMLCII signals in day 7 WT and *PAX3*^−/−^ rosettes. Left to right: VINCULIN-positive cluster area; mean VINCULIN fluorescence intensity within clusters; pMLCII fluorescence intensity within VINCULIN-positive clusters. Colored dots, individual clusters; gray dots, per-experiment means (symbol shape indicates clone); bars, mean ± SEM across independent clone/experiment replicates. **(C)** Top, schematic of the dissociation and replating protocol for day 4 spinal cord organoids onto laminin-coated micropatterns. Bottom, representative images of the resulting micropatterned epithelial monolayers after 7 days of differentiation, stained for F-actin (phalloidin) and DAPI; the left panel is a higher-magnification view of the boxed region in the middle panel, highlighting the honeycomb-like organization of F-actin at apical junctions. **(D)** Representative images of the apical surface of day 7 WT and *PAX3*^−/−^ neural progenitors cultured on micropatterns and stained for F-actin (phalloidin) and CDH2. **(E)** CDH2 fluorescence intensity ratio between junction centers and tricellular junctions, and cell-cell junction straightness, in day 7 WT and *PAX3*^−/−^ neural progenitors cultured on micropatterns. Colored dots, individual junctions; gray dots, per-experiment means; bars, mean ± SEM. **(F)** Representative images of the apical surface of day 7 WT and *PAX3*^−/−^ neural progenitors cultured on micropatterns and labeled with SiR-actin, acquired before and 2 s or 20 s after laser ablation. Pink and blue lines mark the position of the tricellular junction at 0 s and 20 s after ablation, respectively. **(G)** Recoil distance over time after laser ablation in day 7 WT and *PAX3*^−/−^ neural progenitors and in WT progenitors treated with blebbistatin for 10 min, all cultured on micropatterns. Thick lines, mean; shaded areas, SEM. **(H)** Initial recoil velocity derived from (G). Dots, individual ablation events; gray dots, per-experiment means; bars, mean ± SEM. **(I)** Schematic of the VINCULIN tension sensor (Vin-TS), showing the low- and high-tension conformations associated with high and low FRET efficiency, respectively. **(J)** Representative images of the apical surface of WT and *PAX3*^−/−^ neural progenitors expressing Vin-TS or VinT12-TS (T12), cultured on micropatterns, with or without blebbistatin treatment (+Bleb). Panels marked with an asterisk display signal intensity; the remaining panels display normalized FRET-index heatmaps. The color scale for the normalized FRET index is shown below the images. **(K)** Normalized FRET index (WT = 1) at apical cell-cell junctions in progenitors as in (J). Dots, individual cell-cell junctions; gray dots, per-experiment means; bars, mean ± SEM. **(L)** Normalized FRET index (WT = 1) within apical VINCULIN-positive clusters in fixed day 7 WT and *PAX3*^−/−^ organoids expressing the indicated VINCULIN tension sensor construct, with or without blebbistatin treatment. Colored dots, individual VINCULIN-positive clusters; gray dots, per-experiment means; bars, mean ± SEM. Mann-Whitney *P* values are shown on the plots; individual data points, sample sizes, and statistical analyses are provided in Data S1.

We next examined phosphorylated myosin light chain II (pMLCII), a marker of activated non-muscle myosin II and actomyosin contractility. In WT rosettes, pMLCII was enriched in discrete puncta beneath the apical membrane and strongly accumulated at VINCULIN-positive junctional clusters (Fig. 4A and fig. S6A). In contrast, *PAX3*-deficient neuroepithelia retained apical pMLCII puncta, but these appeared more spatially dispersed along the apico-basal axis of the lateral membrane (Fig. 4A and fig. S6A). To quantitatively assess the redistribution of pMLCII, we quantified pMLCII fluorescence within VINCULIN-positive regions, as the punctate organization of the signal precluded reliable segmentation of individual puncta. Total pMLCII signal associated with individual junctional clusters was not reduced in *PAX3*^−/−^ organoids (fig. S6B), whereas mean pMLCII fluorescence intensity was reduced, paralleling the decrease observed for VINCULIN (Fig. 4B). Together, the redistribution of VINCULIN and pMLCII suggests an expansion of mechanically loaded junctional domains following *PAX3* loss. Contractile actomyosin assemblies remain associated with AJs but are distributed across a larger membrane area, resulting in a spatial dilution of junctional contractility rather than a global reduction in force generation. Similar alterations in VINCULIN and pMLCII organization were observed in the dorsal neural tube of E8.5 *Pax3*-mutant embryos (fig. S6, C and D), indicating a conserved consequence of *PAX3* loss in neuroepithelia in humans and mice.

To quantitatively probe the mechanical consequences of *PAX3* loss, we established a two-dimensional neuroepithelial model amenable to live imaging and acute mechanical perturbations by dissociating day 4 spinal organoids and replating neural progenitors onto laminin-coated micropatterns (Fig. 4C). By day 7, the resulting neuroepithelial monolayer was composed predominantly of SOX2⁺ and PAX6⁺ neural progenitors, with PAX3 broadly expressed in a mosaic pattern and no detectable SOX10⁺ neural crest cells (fig. S6E). Cells self-organized into a polarized epithelium with an apical surface exposed to the culture medium and a characteristic honeycomb-like network of F-actin-rich apical junctions (Fig. 4, C and D).

*PAX3*-mutant progenitors likewise formed polarized epithelial monolayers within three days of replating, displaying a characteristic apical honeycomb-like architecture comparable to that of WT cultures (Fig. 4, C and D). Accordingly, key parameters of epithelial cell shape and arrangement, including apical cell area, circularity and neighbour number, were indistinguishable between genotypes (fig. S6F), indicating that *PAX3* loss does not substantially disrupt epithelial organization. Yet, analysis of apical CDH2 organization revealed clear differences between WT and *PAX3*-mutant epithelia (Fig. 4, D and E and fig. S6G). Whereas WT monolayers displayed preferential enrichment of CDH2 at tricellular junctions, *PAX3*-mutant epithelia exhibited a more uniform CDH2 distribution, with increased center-to-vertex intensity ratios (Fig. 4E and fig. S6G). This redistribution was accompanied by reduced junctional straightness (Fig. 4E), suggesting that AJ mechanics may be altered following *PAX3* loss in this model as well. Notably, these defects emerged in geometrically constrained epithelial monolayers assembled on defined laminin substrates, where the ECM is externally imposed rather than cell-derived. This indicates that altered cadherin organization is not simply a secondary consequence of the ECM fragmentation and tissue disorganization observed in *PAX3*-mutant organoids.

To test whether *PAX3* loss alters junctional tension, we performed laser ablation of apical cell-cell contacts in SiR-actin-labeled neuroepithelial monolayers (Fig. 4, F to H and fig. S7A). Laser ablation locally releases pre-existing junctional tension, causing adjacent tricellular junctions to recoil, with recoil dynamics reflecting the tension sustained at the junction. In WT epithelia, ablation induced rapid separation of neighboring tricellular junctions, with recoil distances reaching ∼1.7 µm within 25 s and an initial recoil velocity of ∼0.50 µm s⁻¹ (Fig. 4, F to H and fig. S7A). These values are reminiscent of recoil measurements obtained *in vivo* in the caudal neuroepithelium of the E8.5 mouse embryo (*42*). Treatment with blebbistatin, a myosin II ATPase inhibitor, for 10 min reduced the initial recoil velocity to ∼0.23 µm s⁻¹, indicating that this junctional tension depends on actomyosin contractility. *PAX3*-mutant epithelia also exhibited significantly reduced recoil distance (∼1 µm within 25 s) and initial recoil velocity (∼0.37 µm s⁻¹) following ablation, consistent with reduced junctional tension (Fig. 4, F to H and fig. S7A).

To obtain a measure of junctional tension at a molecular scale, we generated WT and *PAX3*-mutant iPSC lines constitutively expressing the VINCULIN tension sensor (Vin-TS), a FRET-based biosensor that reports tension across individual VINCULIN molecules (*43*) (Fig. 4I). We also generated matched lines expressing the Vin^T12^ tension-sensor variant, carrying four substitutions in the vinculin tail domain (D974A, K975A, R976A and R978A) that weaken the autoinhibitory head-tail interaction and favor VINCULIN activation and tension (*44–46*). iPSC lines were differentiated and processed using this micropattern-based assay for live imaging at day 7. FRET measurements were restricted to VINCULIN-positive clusters at the apical cell-cell interface (Fig. 4J). In WT neuroepithelia, a 10-min treatment with the myosin II inhibitor blebbistatin increased FRET to 1.069 (untreated WT set to 1) (Fig. 4, J and K, and fig. S7B), confirming that Vin-TS reports actomyosin-dependent tension within the neuroepithelium. *PAX3*-deficient neuroepithelia expressing Vin-TS exhibited a significant increase in FRET relative to WT controls (1.040 versus 1.000; Fig. 4, J and K, and fig. S7B), approaching the levels observed after acute myosin II inhibition and indicating substantially reduced tension at adherens junctions. By contrast, cells expressing Vin^T12^-TS displayed uniformly low FRET values (∼0.92), consistent with the constitutively open, tensioned VINCULIN conformation promoted by the T12 mutations (*45*). Similar to blebbistatin treatment, Vin^T12^-TS values were indistinguishable between genotypes (Fig. 4, J and K and fig. S7B), further supporting the conclusion that the differences in FRET observed with Vin-TS between these conditions arise from differences in tension. As Vin-TS measurements have been shown to be largely unaffected by fixation (*47*), we quantified Vin-TS FRET in fixed spinal organoids (Fig. 4L and fig. S7C).

Although mean FRET efficiencies were slightly lower in WT organoids (∼0.958) than in live neuroepithelial monolayers, both blebbistatin treatment (∼0.960) and *PAX3* loss (∼0.959) increased FRET relative to WT controls, consistent with reduced tension at AJs (Fig. 4L and fig. S7C).

These findings identify PAX3 as a regulator of neuroepithelial mechanics. By promoting the spatial confinement of force-bearing AJ domains and maintaining tension across individual VINCULIN molecules, PAX3 enables the efficient concentration of actomyosin-generated forces at AJs. *PAX3* loss disrupts force organization at both the molecular and supramolecular levels, leading to a marked reduction in junctional tension.

### *PAX3 sustains upper glycolytic and glucose-derived biosynthetic pathways in spinal* neuroepithelia

To identify mechanisms linking PAX3 activity to AJ organization and neuroepithelial mechanics, we performed bulk RNA-sequencing on day 7 WT, *PAX3*^+/−^ and *PAX3*^−/−^ spinal organoids (Fig. 5A and fig. S8, A and B and Data S2). Global transcriptional alterations were relatively modest (fig. S8, A and B and Data S2). Principal component analysis (PCA) revealed partial genotype-dependent segregation: *PAX3*^−/−^ samples occupied a transcriptional space distinct from WT controls, although substantial variability remained between replicates and one *PAX3*^+/−^ sample clustered with the WT group (fig. S8A). Accordingly, differential expression analysis identified only 119 significantly deregulated genes, most of which also altered in *PAX3*^−/−^ organoids relative to WT and heterozygous samples (fig. S8B and Data S2).

**Fig. 5.**
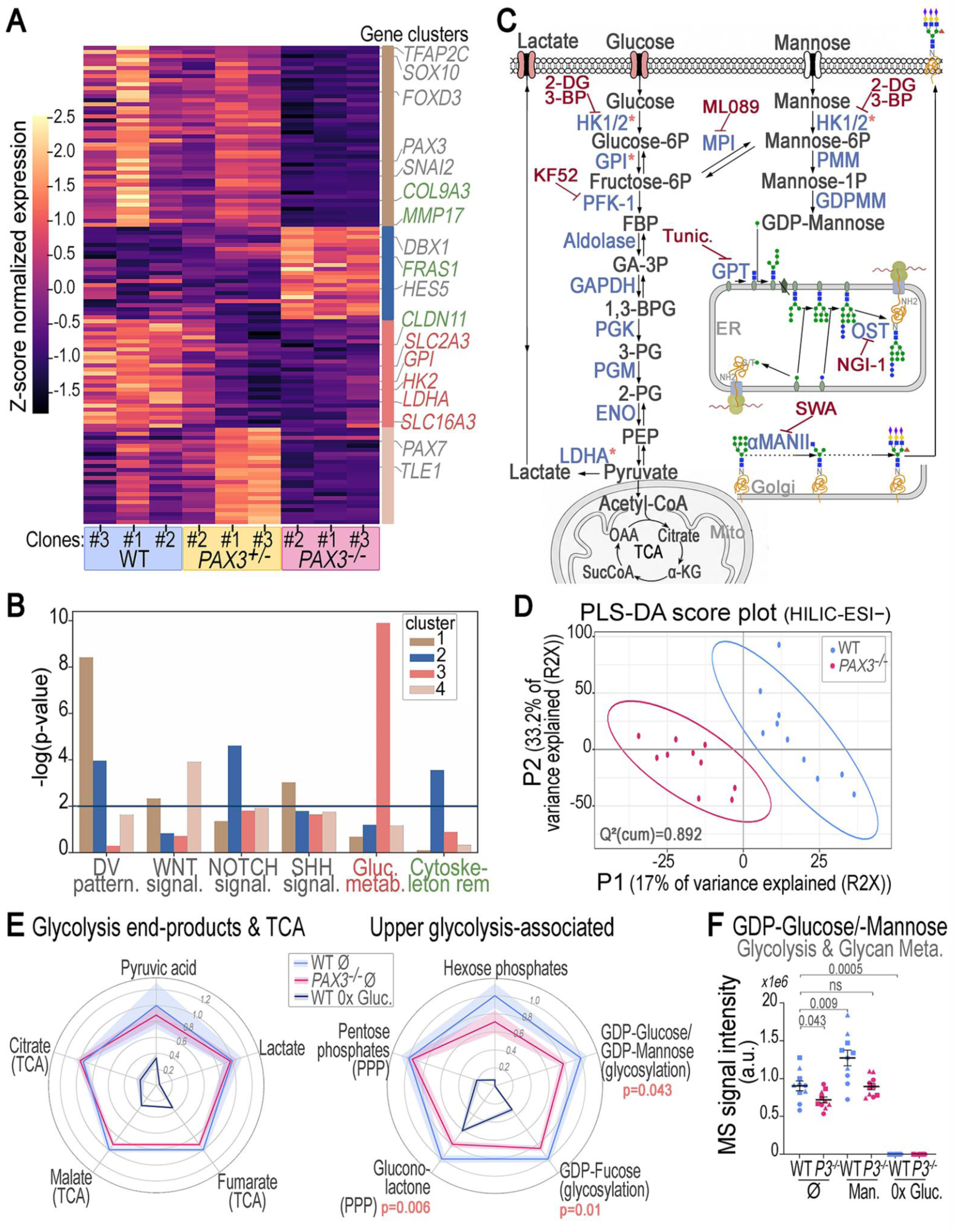
Metabolic dysregulation in *PAX3*-mutant dorsal spinal cord organoids. **(A)** Heatmap of differentially expressed genes identified by RNA sequencing (RNA-seq) of day 7 WT, *PAX3*^+/−^, and *PAX3*^−/−^ organoids (*n* = 3 clones per genotype; dataset in Data S2), revealing four gene clusters with distinct expression dynamics across genotypes. Genes highlighted in gray, green, and pink correspond to dorsoventral patterning genes, regulators of epithelial state, and regulators of glycolysis, respectively. **(B)** Pathway enrichment analysis of the gene clusters identified in (A), highlighting gene sets associated with dorsoventral patterning and patterning-related signaling pathways, epithelial remodeling, and glucose metabolism (glycolysis, TCA cycle, gluconeogenesis). Gene clusters are color-coded as in (A). **(C)** Schematic of the glycolysis and mannose metabolism pathways, indicating the enzymes (blue) and metabolites (black) analyzed in this study and the sites of action of the pharmacological inhibitors used (burgundy). Glucose and lactate transporters highlighted in pink, and enzymes marked with pink stars, are encoded by genes downregulated upon loss of one or both *PAX3* alleles. Abbreviations: HK1/2, hexokinase 1/2; GPI, glucose-6-phosphate isomerase; PFK-1, phosphofructokinase-1; FBP, fructose 1,6-bisphosphate; 2PG, 2- phosphoglyceric acid; GA-3P, glyceraldehyde-3-phosphate; 1,3-BPG, 1,3-bisphosphoglycerate; 3PG, 3-phosphoglycerate; GAPDH, glyceraldehyde-3-phosphate dehydrogenase; PGK, phosphoglycerate kinase; PGM, phosphoglycerate mutase; ENO, enolase; PEP, phosphoenolpyruvate; LDHA, lactate dehydrogenase A; TCA, tricarboxylic acid; OAA, oxaloacetate; αKG, α-ketoglutarate; MPI, mannose phosphate isomerase; PMM, phosphomannomutase; GDP-Man, GDP-mannose; GPT, UDP-*N*-acetylglucosamine phosphotransferase; OST, oligosaccharyltransferase; αManII, α-mannosidase II. Inhibitors: 2-DG, 2-deoxy-D-glucose (hexokinase inhibitor); 3-BP, 3-bromopyruvate (HK1/2 inhibitor); KF52, PFK-1 inhibitor; ML089, MPI inhibitor; NGI-1, OST inhibitor; SWA, swainsonine (αManII inhibitor). **(D)** Partial least squares discriminant analysis (PLS-DA) score plot of HILIC-ESI− metabolomic profiles from day 7 WT and *PAX3*^−/−^ organoids. **(E)** Polar bar diagrams of the relative mass-spectrometric intensity of the indicated metabolites in day 7 WT (blue) and *PAX3*^−/−^ (pink) organoids, and in WT organoids cultured in glucose-free medium (dark blue). Each radial segment corresponds to one metabolite. For each metabolite, intensities are normalized to the maximum across the three conditions (maximum = 1). **(F)** Relative mass-spectrometric intensity of the indicated metabolites in day 7 WT and *PAX3*^−/−^ organoids. Dots, individual organoids; bars, mean ± SEM. For (E) and (F), Mann-Whitney *P* values are shown for comparisons between *PAX3*^−/−^ and WT organoids cultured under basal conditions; sample sizes and statistical analyses are provided in Data S3.

To identify transcriptional programs associated with the morphogenetic phenotype, we clustered differentially expressed genes across the allelic series, revealing four major expression patterns (Fig. 5A and Data S2). One cluster (cluster 2) comprised genes upregulated following *PAX3* loss, including its known direct target *DBX1* (*23, 24*) and a small number of additional patterning-associated transcripts (Fig. 5, A and B, and Data S2). As neural tube closure is rescued by the transcriptional activator PAX3-FOXO1 (*29*), we focused on genes positively regulated by PAX3. Among PAX3-activated genes, cluster 1 was enriched for neural crest-associated transcripts and was selectively reduced in *PAX3*^−/−^ organoids, consistent with the neural crest specification defects described above, whereas cluster 4 lacked significant functional enrichment (Fig. 5, A and B and Data S2). Cluster 3 exhibited a reduction in expression positively correlated with PAX3 dosage and was reduced in the subset of *PAX3*^+/−^ clones displaying the most severe epithelial defects (Fig. 5A), suggesting a closer association with the morphogenetic phenotype. Pathway enrichment analysis identified glucose metabolism (glycolysis, TCA cycle, and gluconeogenesis) as the most significantly enriched biological process within this cluster (Fig. 5B). Downregulated genes included multiple regulators of glucose uptake and upper glycolysis, including SLC2-family glucose transporters (*SLC2A1, SLC2A3*), *HK2*, *GPI*, *LDHA*, as well as the lactate transporter *SLC16A3* (MCT4) (Fig. 5, B and C and fig. S8C). To test whether this holds beyond human cells, we examined our previously published RNA-seq dataset from *Pax3*-deficient mouse spinal organoids (*24*). Glycolytic regulators, including *Hk2* and *Ldha*, were similarly reduced (fig. S8E), indicating that PAX3-dependent maintenance of glycolytic gene expression is conserved between human and mouse. Notably, this glycolytic signature could not be readily explained by altered activity of established regulators of glycolytic gene expression (*48–53*). Wnt, FGF-RAS, PI3K-AKT-mTOR, MYC and hypoxia-responsive pathway activities were unchanged at the transcriptional level in *PAX3*-mutant organoids (fig. S8D).

To determine whether the reduced expression of glycolytic regulators in *PAX3*-mutant organoids was accompanied by metabolic alterations, we performed untargeted metabolomic profiling of day 7 WT and *PAX3*^−/−^ organoids cultured under standard conditions or glucose deprivation (0× glucose). To broaden metabolite coverage, samples were analyzed by two complementary liquid chromatography-tandem mass spectrometry (LC-MS/MS) methods, reversed-phase (C18) and hydrophilic interaction (HILIC) chromatography (Fig. 5, C to E, and fig. S9; Data S3). A total of 10 biological replicates per condition, derived from three independent iPSC clones per genotype, were analyzed. Metabolite annotation identified 129 metabolites that were robustly detected across conditions (Data S3). PLS-DA revealed reproducible metabolic differences between WT and *PAX3*^−/−^ organoids (Fig. 5D and fig. S9B; Data S3). Thirty-three metabolites differed significantly between genotypes and accounted for much of the observed separation (fig. S9B), with 16 decreased and 17 increased in PAX3-mutant organoids. Irrespective of the direction of change in the mutant, 27 of these 33 metabolites were reduced under glucose deprivation (0× glucose) in both genotypes (Fig. 5, E and F and fig. S9, C to E), identifying them as glucose-dependent metabolites. Thus, PAX3 loss preferentially affects metabolites whose abundance depends on glucose availability.

Differentially abundant metabolites mapped to several metabolic pathways (Fig. 5E and fig. S9D). Among these, nucleotide metabolism was prominently remodeled. The purine-derived metabolites xanthine and O-methylinosine were increased, whereas the *de novo* pyrimidine intermediates orotic acid and orotidine were reduced, contrasting with increased levels of the pyrimidine deoxynucleosides thymidine, thymidine monophosphate, and deoxycytidine (Fig. 5E and fig. S9, D and E). This pattern is compatible with reduced *de novo* pyrimidine biosynthesis alongside increased nucleoside salvage or turnover. In addition, several metabolites linked to NAD⁺ metabolism and mitochondrial function, including NAD⁺, cyclic ADP-ribose, α-ketoglutarate, succinate, and several acylcarnitines (carriers of fatty acids into mitochondria), were reduced in *PAX3*-mutant organoids (fig. S9D). By contrast, the glycolytic end products pyruvate and lactate, together with most other tricarboxylic acid (TCA) cycle intermediates such as citrate and malate, remained comparatively preserved, arguing against a major defect in central carbon and energy metabolism (Fig. 5E).

The most coherent signature involved the biosynthetic branches that stem from upper glycolysis. The hexose-phosphates that mark the entry into glycolysis (including glucose-6-phosphate, mannose-6-phosphate, fructose-6-phosphate; detected as a single feature) showed a downward trend in *PAX3*-mutant organoids, although this did not reach statistical significance (Fig. 5E). Two branches fed by these intermediates were more clearly affected. First, glucolactone and the pentose phosphates (including ribose-5-phosphate, ribulose-5-phosphate, and/or xylulose-5-phosphate), products of the pentose phosphate pathway, were significantly reduced, consistent with a lower supply of the sugar precursors used to build nucleotides (Fig. 5E). Second, the activated sugar donors GDP-mannose/GDP-glucose and GDP-fucose were depleted, indicating impaired production of the building blocks required for glycan (sugar-chain) biosynthesis (Fig. 5, C, E, and F).

These transcriptomic and metabolomic data indicate that *PAX3* loss selectively affects glucose-derived biosynthetic pathways rather than energy metabolism. Nucleotide metabolism was remodeled and a subset of NAD⁺ and mitochondrial-carbon-related metabolites was reduced, but the most coherent alterations converged on the upper glycolytic, pentose phosphate, and nucleotide-sugar pathways.

### *PAX3 controls adherens junction organization and mechanics through upper glycolysis and* mannose metabolism

Given the reduction in mannose-derived nucleotide sugars and the established role of glycosylation in regulating cadherin trafficking, stability and adhesive function (*54–56*), we next investigated whether upper glycolytic and mannose-dependent biosynthetic pathways contribute functionally to AJ organization and mechanics. To this end, we pharmacologically perturbed distinct metabolic nodes spanning upper glycolysis, mannose metabolism and glycan biosynthesis in WT and *PAX3*^−/−^ organoids and quantified apical CDH2 and F-actin organization within neuroepithelial rosettes (Fig. 6, A and B and fig. S10).

**Fig. 6.**
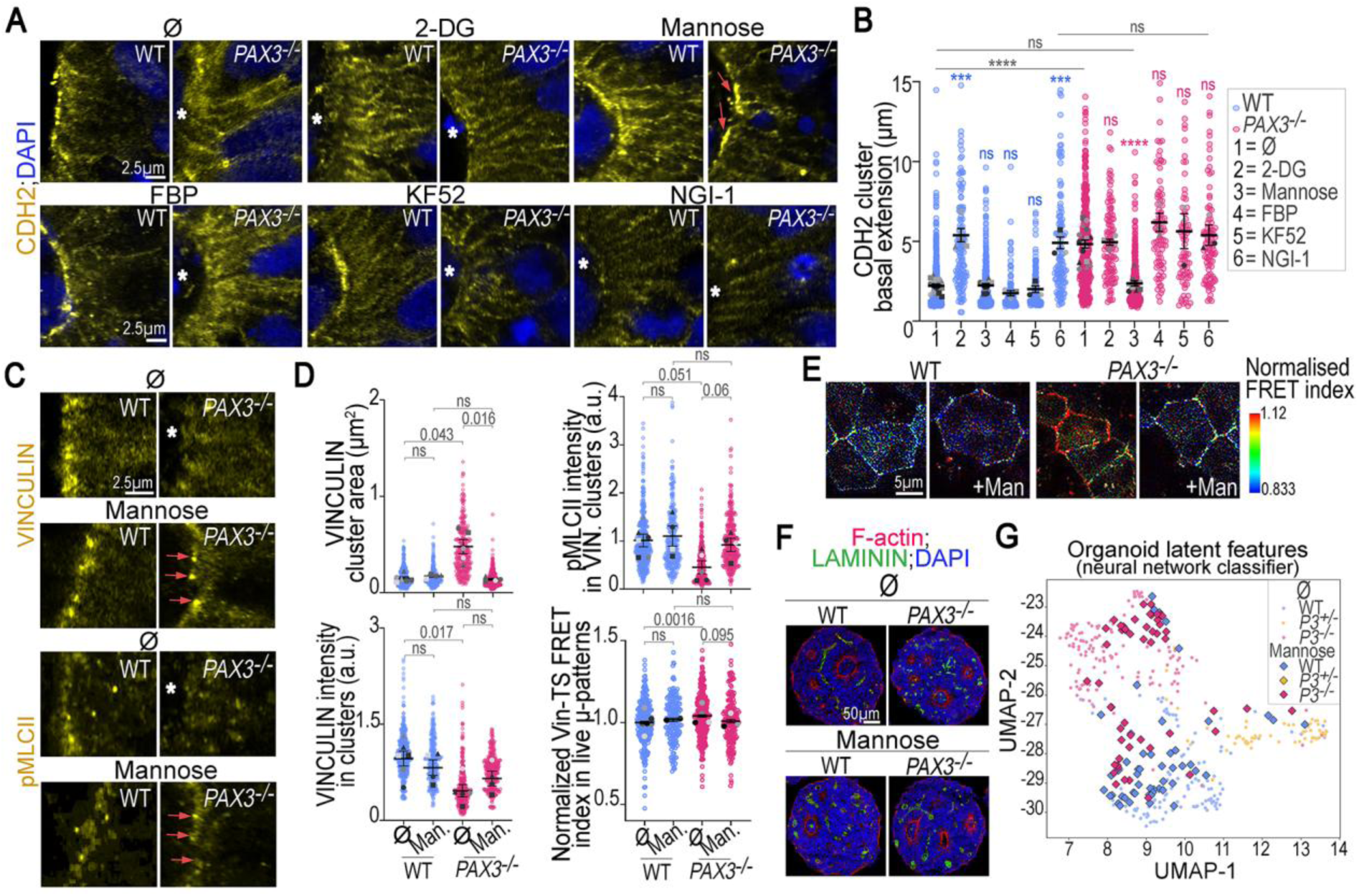
Mannose-mediated restoration of adherens junction clustering, mechanics, and epithelial organization. **(A)** Representative high-magnification images of the apical surface of day 7 WT and *PAX3*^−/−^ organoid rosettes stained for CDH2 and DAPI after treatment with the indicated modulators of glycolysis and mannose metabolism. Asterisks, regions lacking apical CDH2 clusters; pink arrows, rescue of apical CDH2 clustering in *PAX3*^−/−^ organoids after mannose treatment. **(B)** Apical CDH2 cluster extension along the apico-basal axis in organoids as in (A). Colored dots, individual clusters; gray dots, per-experiment means (symbol shape indicates clone); bars, mean ± SEM across independent clone/experiment replicates. For clarity, only selected pairwise comparisons are shown: blue, treated versus untreated WT; gray, WT versus *PAX3*^−/−^; pink, untreated versus treated *PAX3*^−/−^. ns, not significant; ***, P < 0.001; ****, P < 0.0001 (Mann-Whitney test). **(C)** Representative high-magnification images of the apical surface of day 7 WT and *PAX3*^−/−^ organoid rosettes treated with or without mannose and stained for VINCULIN and pMLCII. Asterisks, regions lacking apical VINCULIN or pMLCII clusters; pink arrows, restoration of apical VINCULIN and pMLCII clustering in *PAX3*^−/−^ organoids after mannose treatment. **(D)** Quantifications in day 7 WT and *PAX3*^−/−^ organoids treated with or without mannose: VINCULIN-positive cluster area (top left), mean VINCULIN fluorescence intensity (bottom left), and mean pMLCII fluorescence intensity within VINCULIN-positive clusters (top right); and normalized FRET index at apical cell-cell junctions (bottom right) in day 7 WT and *PAX3*^−/−^ neural progenitors expressing Vin-TS and cultured on micropatterns, treated with or without mannose. Colored dots, individual clusters (top left, bottom left, and top right) or individual cell-cell junctions (bottom right); gray dots, per-experiment means (symbol shape indicates clone; top left, bottom left, and top right); bars, mean ± SEM. **(E)** Representative normalized FRET index heatmaps of apical cell-cell junctions in WT and *PAX3*^−/−^ neural progenitors expressing Vin-TS and cultured on micropatterns, treated with or without mannose. The color scale for the normalized FRET index is shown below the images. **(F)** Representative images of day 7 WT and *PAX3*^−/−^ organoids treated with or without mannose and stained for LAMININ, F-actin (phalloidin), and DAPI. The mannose-treated *PAX3*^−/−^ organoid shown is representative of the rescue phenotype and was classified as WT by the ResNet-50 classifier. **(G)** UMAP projection of latent features from the ResNet-50 classifier trained on images of untreated day 7 WT, *PAX3*^+/−^, and *PAX3*^−/−^ organoids stained for LAMININ, F-actin, and DAPI. Each point, one organoid; circles, untreated organoids; diamonds, mannose-treated organoids projected into the same latent space. Mann-Whitney *P* values are shown on the plots; individual data points, sample sizes, and statistical analyses are provided in Data S1.

We first examined the consequences of limiting glucose availability and its early metabolic utilization. To this end, organoids were cultured in glucose-free medium, in galactose-containing medium, which slows the conversion of sugar-derived carbon into glycolytic intermediates and increases reliance on oxidative metabolism (*57*), or treated with the glucose analogue 2-deoxyglucose (2-DG) or the hexokinase inhibitor 3-bromopyruvate (3-BP), both of which disrupt glucose utilization at or immediately downstream of the hexokinase step. In all cases, WT organoids developed broadened apical CDH2 and F-actin domains that closely resembled the phenotype of *PAX3*-mutant neuroepithelia (Fig. 6, A and B and fig. S10). By contrast, none of these perturbations further exacerbated the phenotype of *PAX3*^−/−^ organoids, suggesting that PAX3 and glucose-dependent metabolic pathways converge on a common mechanism controlling AJ organization. Conversely, we investigated whether supplementing key metabolites could rescue junctional organization in *PAX3*-mutant organoids. Supplementation with excess glucose produced only a modest and variable improvement in apical CDH2 and F-actin organization (Fig. 6, A and B and fig. S10). By contrast, mannose supplementation substantially restored apical clustering of both proteins, while GDP-mannose produced an even more pronounced rescue (Fig. 6, A and B and fig. S10).

Because mannose-6-phosphate can be converted to fructose-6-phosphate through mannose phosphate isomerase (MPI) and thereby re-enter glycolysis, we next asked whether the rescue could reflect restoration of downstream glycolytic metabolism rather than mannose-dependent biosynthetic pathways. Neither fructose-1,6-bisphosphate (FBP) nor pyruvate rescued *PAX3*-mutant organoids (Fig. 6, A and B and fig. S10). Unexpectedly, pyruvate supplementation induced mutant-like CDH2 and F-actin defects in WT tissues (Fig. 6, A and B and fig. S10). As exogenous pyruvate can promote mitochondrial respiration and reduce cellular reliance on upper glycolysis, this observation is consistent with a requirement for glucose-derived metabolites generated upstream of pyruvate production (*58*). Furthermore, inhibition of glycolytic flux downstream of fructose-6-phosphate using the PFK1 inhibitor KF52 had little effect on apical CDH2 or F-actin organization (Fig. 6, A and B and fig. S10). Together, these findings indicate that metabolic processes operating downstream of the fructose-6-phosphate/PFK1 step are largely dispensable for AJ organization and instead point to a requirement for pathways fed upper glycolysis. Further arguing against broad metabolic remodeling as the basis for rescue, metabolomic profiling revealed that mannose supplementation induced remarkably limited changes in both WT and *PAX3*-mutant organoids, with only a small number of metabolites significantly altered (Fig. 5F and fig. S9E and Data S3). Notably, GDP-mannose and GDP-fucose were among the few metabolites increased following treatment, suggesting that exogenous mannose primarily restores nucleotide-sugar metabolism rather than globally rewiring cellular metabolism (Fig. 5F and Data S3).

We therefore asked whether mannose exerts its effects through glycan biosynthesis. Inhibition of mannose phosphate isomerase (MPI, which interconverts fructose-6-phosphate and mannose-6-phosphate) with ML089 phenocopied *PAX3* loss in WT organoids (Fig. 6, A and B, and fig. S10). Likewise, perturbation of N-linked glycosylation at multiple stages, including inhibition of lipid-linked oligosaccharide synthesis (tunicamycin), oligosaccharyltransferase-mediated glycan transfer to nascent proteins (NGI-1), or Golgi α-mannosidase II-dependent glycan maturation (swainsonine), each induced basal spreading of CDH2 and F-actin domains resembling that observed after *PAX3* loss (Figs. 5C and 6, A and B, and fig. S10). These results identify mannose-dependent glycan biosynthesis as a critical downstream effector of PAX3-dependent AJ organization.

The restoration of apical CDH2 and F-actin clustering by mannose supplementation prompted us to determine whether mannose metabolism also rescues the mechanical defects associated with *PAX3* loss. We therefore examined force-bearing AJ assemblies by quantifying VINCULIN organization, pMLCII recruitment and tension across the VINCULIN tension sensor (Vin-TS) in both organoids and neuroepithelial micropattern cultures (Fig. 6, C to E). Strikingly, mannose supplementation restored all three parameters. In *PAX3*-mutant neuroepithelia, mannose restored compact apical VINCULIN clusters resembling those observed in WT controls and increased junction-associated pMLCII levels within these force-bearing domains (Fig. 6, C and D). Consistent with restoration of junctional contractility, mannose treatment also reduced Vin-TS FRET efficiency in both micropattern cultures and fixed organoids, approaching values observed in WT tissues and indicating increased tension across individual VINCULIN molecules (Fig. 6, D and E and fig. S11, A and B). Notably, this rescue occurred without restoring the expression of upper glycolytic genes, indicating that mannose acts downstream of the PAX3-dependent transcriptional defects (fig. S8C). Mannose supplementation additionally increased progenitor proliferation, as reflected by an increased proportion of phospho-histone H3 (PH3)^+^ mitotic cells in both WT and *PAX3*-mutant organoids, but did not rescue cell fate specification defects, as evidenced by the persistent loss of SOX10^+^ neural crest cells and increased DBX1 expression in *PAX3*-mutant organoids (figs. S8C and S11C). Thus, restoration of mannose-dependent metabolism is sufficient not only to rescue AJ organization and mechanics, but also to partially reverse the proliferative defects associated with *PAX3* loss.

Finally, to determine whether restoration of AJ organization and mechanics was accompanied by recovery of higher-order epithelial architecture, we analyzed histological sections stained for LAMININ and F-actin using the ResNet-50 classifier described above (Fig. 6, F and G). As expected, mannose-treated WT organoids clustered with untreated WT controls. In contrast, mannose-treated *PAX3*^−/−^ organoids shifted away from the mutant morphospace toward the WT cluster, and half of them were classified as WT by the classifier. This partial rescue was further supported by silhouette analysis of the UMAP embedding, which quantifies how distinctly the two genotypes separate in feature space: mannose treatment progressively reduced the separation between WT and *PAX3*^−/−^ organoids (fig. S11D). These findings indicate that mannose supplementation partially restores the global neuroepithelial architecture disrupted by *PAX3* loss.

Collectively, these results establish a mechanistic link between the PAX3-dependent metabolic state and neuroepithelial mechanics. By promoting upper glycolytic and mannose-dependent biosynthetic pathways, PAX3 sustains force-bearing adherens junctions, junctional tension, and epithelial organization; restoring mannose metabolism is sufficient to reverse the major structural and mechanical defects caused by *PAX3* loss. These findings identify glycosylation pathways as critical effectors of PAX3-driven neuroepithelial morphogenesis.

## Discussion

Metabolic transitions accompany embryonic development, yet how they are regulated, and which developmental functions they serve, remain largely unknown (*59*). Our study addresses both questions in the context of spinal neurulation: we identify a developmental transcription factor, PAX3, as the regulator of a glycolytic program, and define its output as mechanical rather than fate-determining (*48*, *53*, *60–62*).

In both human and mouse organoids, PAX3 sustains the expression of several genes encoding rate-limiting enzymes of glucose uptake and upper glycolysis (*63*), and it does so independently of the canonical developmental regulators of glycolysis, FGF/Wnt, MYC, HIF1α and PI3K-AKT-mTOR, whose transcriptomic signatures remained largely preserved in *PAX3*-deficient organoids (*49–53*, *60*) (Fig. 5A and fig. S8, D and E). This regulation is very likely to operate *in vivo*: published measurements of glucose uptake, glucose utilization and glycolytic gene expression in mouse and chick embryos reveal a posterior-to-anterior glycolytic gradient not only in the mesoderm but also in the neuroepithelium (*48*, *50*), where it extends further anteriorly, spanning the open neural plate through to the site of closure and coinciding with the PAX3 expression domain. Consistent with a specific effect on the rate-limiting steps of upper glycolysis, *PAX3* loss reduced metabolites of the two biosynthetic branches that diverge from it, namely the pentose phosphate and mannose pathways (Fig. 5, D and E). Strikingly, the glycolytic end products, pyruvate and lactate, were unchanged. One possible explanation is that rapidly proliferating neural progenitors, with their high energetic demand, preferentially maintain carbon flux toward pyruvate to meet ATP and biomass requirements, at the expense of the anabolic branches diverging at glucose-6-phosphate and fructose-6-phosphate (*58*, *59*). Critically, our functional data single out mannose-dependent glycosylation as the biosynthetic branch whose depletion underlies the neuroepithelial mechanical phenotype (Fig. 6, A to E).

This finding bears on a long-standing puzzle in PAX3 biology: the folate responsiveness of *Pax3/Splotch* neural tube defects (*15*, *30–32*). Consistent with previous studies in *Pax3/Splotch* embryos (*15*, *30*, *31*), *PAX3* loss in human spinal neuroepithelial cells is associated with a metabolic signature of impaired *de novo* purine and pyrimidine biosynthesis, processes that rely on folate-derived one-carbon units (Fig. 5E). Mannose supplementation rescued AJ organization and mechanics without restoring this signature (fig. S9E and Data S3), indicating that restoration of junctional mechanics does not require restoration of nucleotide biosynthesis. These findings suggest two non-mutually exclusive possibilities. PAX3 may coordinate distinct biosynthetic pathways supporting complementary processes, with folate-dependent nucleotide synthesis sustaining proliferation (*15*) and mannose-dependent glycosylation promoting junction assembly. Alternatively, these pathways may be metabolically coupled: because GDP-mannose synthesis requires guanine nucleotides, impaired *de novo* purine biosynthesis could limit GDP-mannose production and thereby contribute to the glycosylation defect. Beyond mechanism, these findings broaden the metabolic landscape of neural tube defects beyond folate-nucleotide metabolism, identifying protein glycosylation as an additional metabolic axis that could influence neural tube closure and contribute to the gene-environment interactions underlying these disorders (*6*, *32*).

How is this biosynthetic program ultimately translated into mechanical force? Among the glycoproteins that organise epithelial morphogenesis (*64*), CDH2 is one of the likely effectors. In *PAX3*-deficient neuroepithelia, the adhesive function of CDH2 is largely functional: we do not observe the loss of epithelial integrity seen in CDH2 deficient neuroepithelia (*65*, *66*) (Fig. 2C). However, CDH2 clusters are less confined toward the apical membrane domain. This spatial dilution of CDH2 clusters is accompanied by a reduction in junctional tension, measured at the cell scale by a slower recoil after laser ablation and, at the molecular scale, by a FRET-based vinculin tension sensor (Fig. 4, F to L). The sensor indicates that the mean molecular tension is as low as that measured after acute inhibition of myosin II activity with blebbistatin (Fig. 4, K and L). Yet, the level of contractile myosin (pMLCII) is preserved in *PAX3*^−/−^ neuroepithelia, but its organization is altered and more diffuse (Fig. 4, A and B and fig. S6, A and B). The primary defect would therefore lie in the transmission of contractility rather than in the upstream signal generating contractile forces (*67*). More importantly, supplementing *PAX3*-mutant organoids with mannose is sufficient to restore CDH2 clustering, vinculin and pMLCII organization, and the tension across the vinculin sensor (Fig. 6, A to D). Although we cannot hierarchize CDH2 clustering and junctional tension causally, they are co-regulated in our system as in many others (*39*, *40*). Our data show that glycosylation lies upstream of this force-bearing state and of both CDH2 clustering and junctional tension (Fig. 6, A to D and fig. S10, A to C). This is reinforced by our finding that blocking N-glycosylation at multiple independent steps, from the formation of simple glycans in the ER to that of complex glycans in the Golgi apparatus, disperses apical CDH2 and F-actin in WT neuroepithelia (Fig. 6, A and B and fig. S10, A to C).

How the glycan structures of CDH2 govern this force-bearing state remains to be defined. Studies of CDH2 and CDH1 at both the molecular and cellular levels show that the amount of glycans and the nature of the glycan trees modulate cadherin adhesion strength and binding kinetics, with simple, highly mannosylated trees favoring these properties (*55*, *56*, *68*). We hypothesise that a similar principle may apply to the force-bearing clustering that we measure, although this remains to be tested directly. We cannot rule out the involvement of other glycosylated targets. In the mesoderm, where mannose can likewise rescue the mesoderm specification and axial elongation of gastruloids impaired by glycolytic blockade, N-glycosylation of secreted Wnt-pathway modulators such as Sfrp3 has been proposed to be involved (*62*). Because Wnt signaling is transcriptionally preserved in the *PAX3*-deficient neuroepithelium, we favor the view that the glycosylated effectors vary between tissues, while glycosylation itself constitutes a unifying metabolic input into the caudal morphogenesis of the embryo.

Finally, restoring mannose metabolism rescued not only junctional tension but also, at least partially, the higher-order neuroepithelial architecture disrupted by *PAX3* loss (Fig. 6, F and G). This links the spatial organization of junctional forces to tissue-scale morphogenesis, and is consistent with *in vivo* evidence that a properly organized apical actomyosin and adherens-junction network is required for neural fold morphogenesis and neural tube closure, since dysregulating this organization impairs closure (*69*, *70*). Together, our results define a pathway in which a developmental transcription factor, through an upper-glycolytic and mannose-dependent glycosylation program, confines contractile force at adherens junctions and thereby endows the human spinal neuroepithelium with the mechanical competence required for its morphogenesis.

## Acknowledgments

We sincerely thank Pierre-Luc Bardet for his insightful input, thoughtful discussions and continued support throughout the years of this project. His influence was instrumental in shaping this work. We thank Gaël Simon, Mariia Balatskaia, Benoît Sorre, Julie Stoufflet, René-Marc Mège, Cyril Hanus, Jean-Marc Verbavatz, Valérie Doye, Auguste Genovesio, and Valérie Mezger for valuable discussions and support. We also acknowledge the ImagoSeine core facility of the Institut Jacques Monod, member of the France BioImaging infrastructure (https://ror.org/01y7vt929) supported by the French National Research Agency (ANR-24-INBS-0005 FBI BIOGEN) and GIS-IBiSA, as well as the EnSCORE platform for assistance with iPSC handling. Finally, we thank James Briscoe and Despina Stamataki for critical comments on the manuscript. During the preparation of this work, the authors used ChatGPT (OpenAI) and Claude (Anthropic) to assist with language editing, text condensation, and the identification of potential interactions within metabolic pathways. Following the use of these tools, the authors critically reviewed, revised, and validated all generated content and take full responsibility for the accuracy, integrity, and originality of the published work.

## Author contributions

Conceptualization: C.M., V.R., N.B., S.N., S.L.

Methodology: C.M., X.B., H.L., M.B.

Software: A.B., S.A., H.L., C.M.

Validation: K.B., M.B., C.M., V.R., S.A., L.D., B.E.

Formal analysis: C.M., H.L., S.A., L.D., A.B., F.C., C.C., E.C.V., V.R.

Investigation: C.M., L.D., K.B., M.B., S.A., B.E., N.V., F.C., C.C., E.C.V., S.B., A.M., C.D., V.R.

Resources: V.R., F.R., S.N., N.B.

Data curation: C.M., L.D., F.C.

Writing - original draft: C.M., V.R.

Writing - review & editing: N.B., S.N., S.L.

Visualization: C.M., V.R.

Supervision: V.R., N.B.

Funding acquisition: V.R., N.B., S.N., V.D., F.R.

## Diversity, equity, ethics, and inclusion

Our author team is gender-balanced and brings together researchers at multiple career stages, from PhD students to senior investigators, across several disciplines (developmental biology, biophysics, metabolomics, and computational image analysis) and institutions. We are committed to equitable and inclusive research practices and strove to reflect this in team composition, authorship, and mentoring of early-career scientists.

## Data, code, and materials availability

Data, code, and materials availability: RNA-seq data have been deposited in the Gene Expression Omnibus (GEO) under accession number GSE341658. Mass spectrometry-based metabolomics data have been deposited in the MassIVE repository (https://massive.ucsd.edu) under accession number MassIVE MSV000102730 (DOI: 10.25345/C58W38H1M). Analysis code, including image analysis and AI-based analysis pipelines, is available on GitHub (https://github.com/ribeslab/Mirdass-et-al), ensuring long-term accessibility. Newly generated iPSC lines and other materials are available from the corresponding authors upon request, subject to applicable material transfer agreements. All other data supporting the findings of this study are available in the manuscript or the supplementary materials.

## Supplementary Materials

Materials and Methods

Figs. S1 to S11

Tables S1 to S3

References (*1–14*)

Data S1 to S3

## References

1. D. Gilmour, M. Rembold, M. Leptin, From morphogen to morphogenesis and back. Nature 541, 311–320 (2017).

2. C. Collinet, T. Lecuit, Programmed and self-organized flow of information during morphogenesis. Nat. Rev. Mol. Cell Biol. 22, 245–265 (2021).

3. O. Campàs, A toolbox to explore the mechanics of living embryonic tissues. Semin. Cell Dev. Biol. 55, 119–130 (2016).

4. F. Spitz, E. E. M. Furlong, Transcription factors: from enhancer binding to developmental control. Nat. Rev. Genet. 13, 613–626 (2012).

5. E. Nikolopoulou, G. L. Galea, A. Rolo, N. D. E. Greene, A. J. Copp, Neural tube closure: cellular, molecular and biomechanical mechanisms. Development 144, 552–566 (2017).

6. S. Lee, J. G. Gleeson, Closing in on Mechanisms of Open Neural Tube Defects. Trends Neurosci. 43, 519–532 (2020).

7. K. S. Au, L. Hebert, P. Hillman, C. Baker, M. R. Brown, D.-K. Kim, K. Soldano, M. Garrett, A. Ashley-Koch, S. Lee, J. Gleeson, J. E. Hixson, A. C. Morrison, H. Northrup, Human myelomeningocele risk and ultra-rare deleterious variants in genes associated with cilium, WNT-signaling, ECM, cytoskeleton and cell migration. Sci. Rep. 11, 3639 (2021).

8. A. J. Copp, N. S. Adzick, L. S. Chitty, J. M. Fletcher, G. N. Holmbeck, G. M. Shaw, Spina bifida. Nat. Rev. Dis. Primer 1, 15007 (2015).

9. Y.-J. J. Ha, A. Nisal, I. Tang, C. Lee, I. Jhamb, C. Wallace, R. Howarth, S. Schroeder, K. I. Vong, N. Meave, F. Jiwani, C. Barrows, S. Lee, N. Jiang, A. Patel, K. Bagga, N. Banka, L. Friedman, F. A. Blanco, S. Yu, S. Rhee, H. S. Jeong, I. Plutzer, M. B. Major, B. Benoit, C. Poüs, C. Heffner, Z. Kibar, G. M. Bot, H. Northrup, K. S. Au, M. Strain, A. E. Ashley-Koch, R. H. Finnell, J. T. Le, H. S. Meltzer, C. Araujo, H. R. Machado, R. E. Stevenson, A. Yurrita, S. Mumtaz, A. Ahmed, M. H. Khara, O. M. Mutchinick, J. R. Medina-Bereciartu, F. Hildebrandt, G. Melikishvili, A. I. Marwan, V. Capra, M. M. Noureldeen, A. M. S. Salem, M. Y. Issa, M. S. Zaki, L. Xu, J. E. Lee, D. Shin, A. Alkelai, A. R. Shuldiner, S. F. Kingsmore, S. A. Murray, H. Y. Gee, W. T. Miller, K. F. Tolias, J. B. Wallingford, Spina Bifida Sequencing Consortium, A. E. A. Koch, H. S. Meltzer, J. T. Le, K. S. Au, P. J. Lupo, C. Araújo, T. Magana, C. M. Kolvenbach, S. Shril, Y. Takahashi, H. Salimi-Dafsari, H. W. Phillips, B. Hanak, B. Kara, A. S. Güneş, D. D. Gonda, S. Kirmani, T. Tkemaladze, S. Kim, J. G. Gleeson, The contribution of de novo coding mutations to meningomyelocele. Nature 641, 419–426 (2025).

10. M. D. Goulding, G. Chalepakis, U. Deutsch, J. R. Erselius, P. Gruss, Pax-3, a novel murine DNA binding protein expressed during early neurogenesis. EMBO J. 10, 1135–1147 (1991).

11. A. Kicheva, T. Bollenbach, A. Ribeiro, H. P. Valle, R. Lovell-Badge, V. Episkopou, J. Briscoe, Coordination of progenitor specification and growth in mouse and chick spinal cord. Science 345, 1254927 (2014).

12. O. Sanchez-Ferras, G. Bernas, E. Laberge-Perrault, N. Pilon, Induction and dorsal restriction of Paired-box 3 (Pax3) gene expression in the caudal neuroectoderm is mediated by integration of multiple pathways on a short neural crest enhancer. Biochim. Biophys. Acta BBA - Gene Regul. Mech. 1839, 546–558 (2014).

13. M. Zagorski, Y. Tabata, N. Brandenberg, M. P. Lutolf, G. Tkačik, T. Bollenbach, J. Briscoe, A. Kicheva, Decoding of position in the developing neural tube from antiparallel morphogen gradients. Science 356, 1379–1383 (2017).

14. D. J. Epstein, M. Vekemans, P. Gros, splotch (Sp2H), a mutation affecting development of the mouse neural tube, shows a deletion within the paired homeodomain of Pax-3. Cell 67, 767–774 (1991).

15. S. Sudiwala, A. Palmer, V. Massa, A. J. Burns, L. P. E. Dunlevy, S. C. P. De Castro, D. Savery, K.-Y. Leung, A. J. Copp, N. D. E. Greene, Cellular mechanisms underlying Pax3 related neural tube defects and their prevention by folic acid. Dis. Model. Mech., dmm.042234 (2019).

16. N. D. E. Greene, V. Massa, A. J. Copp, Understanding the causes and prevention of neural tube defects: Insights from the *splotch* mouse model. Birt. Defects Res. A. Clin. Mol. Teratol. 85, 322–330 (2009).

17. G. Chalepakis, M. Goulding, A. Read, T. Strachan, P. Gruss, Molecular basis of splotch and Waardenburg Pax-3 mutations. Proc. Natl. Acad. Sci. 91, 3685–3689 (1994).

18. A. J. Agopian, A. D. Bhalla, E. Boerwinkle, R. H. Finnell, M. L. Grove, J. E. Hixson, L. C. Shimmin, A. Sewda, C. Stuart, Y. Zhong, H. Zhu, L. E. Mitchell, Exon sequencing of *PAX3* and *T* ( *Brachyury* ) in cases with spina bifida: *PAX3* , *T* , AND SPINA BIFIDA. Birt. Defects Res. A. Clin. Mol. Teratol., n/a-n/a (2013).

19. S. Chatkupt, F. A. Hol, Y. Y. Shugart, M. P. Geurds, E. S. Stenroos, M. R. Koenigsberger, B. C. Hamel, W. G. Johnson, E. C. Mariman, Absence of linkage between familial neural tube defects and PAX3 gene. J. Med. Genet. 32, 200–204 (1995).

20. J. Hart, K. Miriyala, Neural tube defects in Waardenburg syndrome: A case report and review of the literature. Am. J. Med. Genet. A. 173, 2472–2477 (2017).

21. F. A. Hol, B. C. Hamel, M. P. Geurds, R. A. Mullaart, F. G. Barr, R. A. Macina, E. C. Mariman, A frameshift mutation in the gene for PAX3 in a girl with spina bifida and mild signs of Waardenburg syndrome. J. Med. Genet. 32, 52–56 (1995).

22. P. Lemay, M.-C. Guyot, É. Tremblay, A. Dionne-Laporte, D. Spiegelman, É. Henrion, O. Diallo, P. De Marco, E. Merello, C. Massicotte, V. Désilets, J. L. Michaud, G. A. Rouleau, V. Capra, Z. Kibar, Loss-of-function de novo mutations play an important role in severe human neural tube defects. J. Med. Genet. 52, 493–497 (2015).

23. C. Gard, G. Gonzalez Curto, Y. E.-M. Frarma, E. Chollet, N. Duval, V. Auzié, F. Auradé, L. Vigier, F. Relaix, A. Pierani, F. Causeret, V. Ribes, Pax3- and Pax7-mediated Dbx1 regulation orchestrates the patterning of intermediate spinal interneurons. Dev. Biol. 432, 24–33 (2017).

24. R. Rondon, T. Hezez, J. Richard Albert, S. Hayashi, B. Drayton-Libotte, G. G. Curto, F. Auradé, E. Balloul, C. Dugast-Darzacq, F. Relaix, P. Gilardi-Hebenstreit, V. Ribes, Dual transcriptional activities of PAX3 and PAX7 spatially encode spinal cell fates through distinct gene networks. PLOS Biol. 23, e3003448 (2025).

25. T. Sato, N. Sasai, Y. Sasai, Neural crest determination by co-activation of *Pax3* and *Zic1* genes in *Xenopus* ectoderm. Development 132, 2355–2363 (2005).

26. J.-L. Plouhinec, D. D. Roche, C. Pegoraro, A. L. Figueiredo, F. Maczkowiak, L. J. Brunet, C. Milet, J.-P. Vert, N. Pollet, R. M. Harland, A. H. Monsoro-Burq, Pax3 and Zic1 trigger the early neural crest gene regulatory network by the direct activation of multiple key neural crest specifiers. Dev. Biol. 386, 461–472 (2014).

27. H. Nakazaki, A. C. Reddy, B. L. Mania-Farnell, Y.-W. Shen, S. Ichi, C. McCabe, D. George, D. G. McLone, T. Tomita, C. S. K. Mayanil, Key basic helix–loop–helix transcription factor genes Hes1 and Ngn2 are regulated by Pax3 during mouse embryonic development. Dev. Biol. 316, 510–523 (2008).

28. S. Ichi, V. Boshnjaku, Y.-W. Shen, B. Mania-Farnell, S. Ahlgren, S. Sapru, N. Mansukhani, D. G. McLone, T. Tomita, C. S. K. Mayanil, Role of Pax3 acetylation in the regulation of *Hes1* and *Neurog2*. Mol. Biol. Cell 22, 503–512 (2011).

29. F. Relaix, M. Polimeni, D. Rocancourt, C. Ponzetto, B. W. Schäfer, M. Buckingham, The transcriptional activator PAX3-FKHR rescues the defects of *Pax3* mutant mice but induces a myogenic gain-of-function phenotype with ligand-independent activation of Met signaling in vivo. Genes Dev. 17, 2950–2965 (2003).

30. A. Fleming, A. J. Copp, Embryonic Folate Metabolism and Mouse Neural Tube Defects. Science 280, 2107–2109 (1998).

31. A. E. Beaudin, E. V. Abarinov, D. M. Noden, C. A. Perry, S. Chu, S. P. Stabler, R. H. Allen, P. J. Stover, Shmt1 and de novo thymidylate biosynthesis underlie folate-responsive neural tube defects in mice. Am. J. Clin. Nutr. 93, 789–798 (2011).

32. K. A. Burren, D. Savery, V. Massa, R. M. Kok, J. M. Scott, H. J. Blom, A. J. Copp, N. D. E. Greene, Gene-environment interactions in the causation of neural tube defects: folate deficiency increases susceptibility conferred by loss of Pax3 function. Hum. Mol. Genet. 17, 3675–3685 (2008).

33. B. J. Wlodarczyk, L. S. Tang, A. Triplett, F. Aleman, R. H. Finnell, Spontaneous neural tube defects in splotch mice supplemented with selected micronutrients. Toxicol. Appl. Pharmacol. 213, 55–63 (2006).

34. N. Duval, C. Vaslin, T. Barata, Y. Frarma, V. Contremoulins, X. Baudin, S. Nédélec, V. Ribes, BMP4 patterns Smad activity and generates stereotyped cell fate organisation in spinal organoids. Development, dev.175430 (2019).

35. M. Saade, E. Martí, Early spinal cord development: from neural tube formation to neurogenesis. Nat. Rev. Neurosci. 26, 195–213 (2025).

36. A. J. Blasky, A. Mangan, R. Prekeris, Polarized Protein Transport and Lumen Formation During Epithelial Tissue Morphogenesis. Annu. Rev. Cell Dev. Biol. 31, 575–591 (2015).

37. C. E. Buckley, D. St Johnston, Apical–basal polarity and the control of epithelial form and function. Nat. Rev. Mol. Cell Biol. 23, 559–577 (2022).

38. C. Mirdass, M. Catala, M. Bocel, S. Nedelec, V. Ribes, Stem cell-derived models of spinal neurulation. Emerg. Top. Life Sci. 7, 423–437 (2023).

39. S. M. Troyanovsky, Adherens junction: the ensemble of specialized cadherin clusters. Trends Cell Biol. 33, 374–387 (2023).

40. O. Campàs, I. Noordstra, A. S. Yap, Adherens junctions as molecular regulators of emergent tissue mechanics. Nat. Rev. Mol. Cell Biol. 25, 252–269 (2024).

41. S. Yonemura, Y. Wada, T. Watanabe, A. Nagafuchi, M. Shibata, α-Catenin as a tension transducer that induces adherens junction development. Nat. Cell Biol. 12, 533–542 (2010).

42. L. Bocanegra-Moreno, A. Singh, E. Hannezo, M. Zagorski, A. Kicheva, Cell cycle dynamics control fluidity of the developing mouse neuroepithelium. Nat. Phys. 19, 1050– 1058 (2023).

43. C. Grashoff, B. D. Hoffman, M. D. Brenner, R. Zhou, M. Parsons, M. T. Yang, M. A. McLean, S. G. Sligar, C. S. Chen, T. Ha, M. A. Schwartz, Measuring mechanical tension across vinculin reveals regulation of focal adhesion dynamics. Nature 466, 263–266 (2010).

44. H. Canever, H. Lachuer, Q. Delaunay, F. Sipieter, N. Audugé, P. P. Girard, N. Borghi, Collective directional memory controls the range of epithelial cell migration. [Preprint] (2026). 10.7554/eLife.110739.1.

45. D. S. Chorev, T. Volberg, A. Livne, M. Eisenstein, B. Martins, Z. Kam, B. M. Jockusch, O. Medalia, M. Sharon, B. Geiger, Conformational states during vinculin unlocking differentially regulate focal adhesion properties. Sci. Rep. 8, 2693 (2018).

46. D. M. Cohen, H. Chen, R. P. Johnson, B. Choudhury, S. W. Craig, Two Distinct Head-Tail Interfaces Cooperate to Suppress Activation of Vinculin by Talin. J. Biol. Chem. 280, 17109–17117 (2005).

47. E. M. Gates, A. S. LaCroix, K. E. Rothenberg, B. D. Hoffman, Improving Quality, Reproducibility, and Usability of FRET-Based Tension Sensors. Cytometry A 95, 201– 213 (2019).

48. V. Bulusu, N. Prior, M. T. Snaebjornsson, A. Kuehne, K. F. Sonnen, J. Kress, F. Stein, C. Schultz, U. Sauer, A. Aulehla, Spatiotemporal Analysis of a Glycolytic Activity Gradient Linked to Mouse Embryo Mesoderm Development. Dev. Cell 40, 331–341.e4 (2017).

49. T. S. Cliff, T. Wu, B. R. Boward, A. Yin, H. Yin, J. N. Glushka, J. H. Prestegaard, S. Dalton, MYC Controls Human Pluripotent Stem Cell Fate Decisions through Regulation of Metabolic Flux. Cell Stem Cell 21, 502–516.e9 (2017).

50. M. Oginuma, P. Moncuquet, F. Xiong, E. Karoly, J. Chal, K. Guevorkian, O. Pourquié, A Gradient of Glycolytic Activity Coordinates FGF and Wnt Signaling during Elongation of the Body Axis in Amniote Embryos. Dev. Cell 40, 342–353.e10 (2017).

51. R. A. Saxton, D. M. Sabatini, mTOR Signaling in Growth, Metabolism, and Disease. Cell 168, 960–976 (2017).

52. G. L. Semenza, HIF-1: upstream and downstream of cancer metabolism. Curr. Opin. Genet. Dev. 20, 51–56 (2010).

53. A. Villaronga-Luque, R. G. Savill, N. López-Anguita, A. Bolondi, S. Garai, S. I. Gassaloglu, R. Rouatbi, K. Schmeisser, A. Poddar, L. Bauer, T. Alves, S. Traikov, J. Rodenfels, T. Chavakis, A. Bulut-Karslioglu, J. V. Veenvliet, Integrated molecular-phenotypic profiling reveals metabolic control of morphological variation in a stem-cell-based embryo model. Cell Stem Cell 32, 759–777.e13 (2025).

54. C.-L. Zhang, C. Moutoussamy, M. Tuffery, A. Varangot, R. Piskorowski, C. Hanus, Core-N-glycans are atypically abundant at the neuronal surface and regulate glutamate receptor signaling. Cell Biology [Preprint] (2024). 10.1101/2024.03.25.586577.

55. M. D. Langer, H. Guo, N. Shashikanth, J. M. Pierce, D. E. Leckband, N-Glycosylation Alters Cadherin-Mediated Intercellular Binding Kinetics. J. Cell Sci., jcs.101147 (2012).

56. H.-B. Guo, H. Johnson, M. Randolph, M. Pierce, Regulation of Homotypic Cell-Cell Adhesion by Branched N-Glycosylation of N-cadherin Extracellular EC2 and EC3 Domains. J. Biol. Chem. 284, 34986–34997 (2009).

57. R. Shiratori, K. Furuichi, M. Yamaguchi, N. Miyazaki, H. Aoki, H. Chibana, K. Ito, S. Aoki, Glycolytic suppression dramatically changes the intracellular metabolic profile of multiple cancer cell lines in a mitochondrial metabolism-dependent manner. Sci. Rep. 9, 18699 (2019).

58. A. R. Diers, K. A. Broniowska, C.-F. Chang, N. Hogg, Pyruvate fuels mitochondrial respiration and proliferation of breast cancer cells: effect of monocarboxylate transporter inhibition. Biochem. J. 444, 561–571 (2012).

59. T. S. Tippetts, M. H. Sieber, A. Solmonson, Beyond energy and growth: the role of metabolism in developmental signaling, cell behavior and diapause. Development 150, dev201610 (2023).

60. M. Oginuma, Y. Harima, O. A. Tarazona, M. Diaz-Cuadros, A. Michaut, T. Ishitani, F. Xiong, O. Pourquié, Intracellular pH controls WNT downstream of glycolysis in amniote embryos. Nature 584, 98–101 (2020).

61. K. S. Stapornwongkul, E. Hahn, P. Poliński, L. Salamó Palau, K. Arató, L. Yao, K. Williamson, N. Gritti, K. Anlas, M. Osuna Lopez, K. R. Patil, I. Heemskerk, M. Ebisuya, V. Trivedi, Glycolytic activity instructs germ layer proportions through regulation of Nodal and Wnt signaling. Cell Stem Cell 32, 744–758.e7 (2025).

62. C. Dingare, D. Cao, J. J. Yang, B. Sozen, B. Steventon, Mannose controls mesoderm specification and symmetry breaking in mouse gastruloids. Dev. Cell 59, 1523–1537.e6 (2024).

63. L. B. Tanner, A. G. Goglia, M. H. Wei, T. Sehgal, L. R. Parsons, J. O. Park, E. White, J. E. Toettcher, J. D. Rabinowitz, Four Key Steps Control Glycolytic Flux in Mammalian Cells. Cell Syst. 7, 49–62.e8 (2018).

64. M. He, X. Zhou, X. Wang, Glycosylation: mechanisms, biological functions and clinical implications. Signal Transduct. Target. Ther. 9, 194 (2024).

65. M. Kadowaki, S. Nakamura, O. Machon, S. Krauss, G. L. Radice, M. Takeichi, N-cadherin mediates cortical organization in the mouse brain. Dev. Biol. 304, 22–33 (2007).

66. S. I. I. Gänzler-Odenthal, C. Redies, Blocking N-Cadherin Function Disrupts the Epithelial Structure of Differentiating Neural Tissue in the Embryonic Chicken Brain. J. Neurosci. 18, 5415–5425 (1998).

67. A. S. Yap, K. Duszyc, V. Viasnoff, Mechanosensing and Mechanotransduction at Cell– Cell Junctions. Cold Spring Harb. Perspect. Biol. 10, a028761 (2018).

68. A. Liwosz, T. Lei, M. A. Kukuruzinska, N-Glycosylation Affects the Molecular Organization and Stability of E-cadherin Junctions. J. Biol. Chem. 281, 23138–23149 (2006).

69. S. Escuin, B. Vernay, D. Savery, C. B. Gurniak, W. Witke, N. D. E. Greene, A. J. Copp, Rho kinase-dependent actin turnover and actomyosin disassembly are necessary for mouse spinal neural tube closure. J. Cell Sci., jcs.164574 (2015).

70. T. Nishimura, H. Honda, M. Takeichi, Planar Cell Polarity Links Axes of Spatial Dynamics in Neural-Tube Closure. Cell 149, 1084–1097 (2012).

