## Supplementary Materials for "Transcriptional coupling of metabolism and cell mechanics controls human neuroepithelium morphogenesis"

**This PDF file includes:**

Materials and Methods
Figs. S1 to S11
Tables S1 to S3
References

**Other Supplementary Materials for this manuscript include the following:**

Data S1 to S3

### Materials and Methods

Human materials and ethics statement

This study used the human induced pluripotent stem cell line WTC-11 (Coriell Institute for Medical Research, catalogue no. GM25256), derived from a healthy adult donor who gave written informed consent for research use. In accordance with the French Public Health Code, the conservation and use of human biological samples for scientific research in our laboratory are covered by CODECOH declaration no. DC-2021-4363 to the French Ministry of Higher Education and Research (articles L.1243-3 and L.1243-4 CSP). All genetically modified derivatives used here (*PAX3* alleles, *Vin-TS* transgenes) were generated from this parental line in our laboratory; no new line was derived from primary human material for this study. The contained use of these genetically modified organisms (the WTC-11 parental line and its *PAX3*-mutant and *Vin-TS*-expressing derivatives) was declared and approved under the French regulations on the contained use of GMOs (dossier DUO no. 12325, French Ministry of Higher Education and Research). The work did not fall under the protocols requiring declaration to the Agence de la biomédecine under the French bioethics law (loi no. 2021-1017; article L.2151-7 CSP).

Human iPSC culture and maintenance

Two human induced pluripotent stem cell (iPSC) lines derived from the WTC-11 background were used in this study: the wild-type (WT) line WTC-11 (male; WTC/AICS-0/GM25256; RRID:CVCL_Y803) and an endogenously tagged β-CATENIN-GFP reporter line (CTNNB1-GFP; AICS-0058-067; RRID:CVCL_VK86), both obtained from the National Institute of General Medical Sciences Human Genetic Cell Repository at the Coriell Institute for Medical Research. WTC-11 was used as the parental line for the generation of the *PAX3*-mutant and *Vin-TS* transgenic lines, and served as the WT control throughout the study unless otherwise stated. The β-CATENIN-GFP reporter line was used specifically for characterizing the timing of PAX3 induction and of the progressive formation of neuroepithelial structures over time (Fig. 1D).

All iPSCs were cultured on hESC-qualified Matrigel-coated plates (Corning; #11553620, Fisher Scientific) in mTeSR Plus medium (#100-0276, STEMCELL Technologies) at 37 °C and 5% $CO_2$, and passaged as small aggregates using Gentle Cell Dissociation Reagent (#100-0485, STEMCELL Technologies).

Generation of *PAX3*-mutant iPSC lines

Human iPSC lines carrying heterozygous or homozygous CRISPR-Cas9-induced frameshift null mutations in *PAX3* were generated from the parental WTC-11 line. crRNA targeting exon 2 of *PAX3* was designed using CRISPOR (http://crispor.tefor.net)(*1*) and selected on the basis of predicted efficiency and specificity. Guide RNA (gRNA) was generated by annealing equimolar Alt-R CRISPR-Cas9 crRNA and Alt-R CRISPR-Cas9 tracrRNA ATTO 550 (both Integrated DNA Technologies, IDT) at 95 °C for 5 min, followed by cooling to room temperature for 20 min. Ribonucleoprotein (RNP) complexes were assembled by incubating 150 pmol of gRNA with 122 pmol of Alt-R S.p. HiFi Cas9 Nuclease V3 (IDT) for 20 min at room temperature. WTC-11 iPSCs at passage 45 were treated with 10 µM of the ROCK inhibitor Y-27632 (Axon Medchem) for 1 h and dissociated into single cells using Gentle Cell Dissociation Reagent. A total of $6 \times 10^5$ cells were resuspended in 30 µL of supplemented P3 buffer (Lonza) and mixed with the RNP complex. Nucleofection was performed using the 4D-Nucleofector X Unit (Lonza) with program CA137. Cells were seeded in 4-well plates (Thermo Scientific) at 100,000 cells/cm² in mTeSR Plus medium supplemented with CloneR2

(STEMCELL Technologies). Twenty-four hours after nucleofection, cells were dissociated as described above and sorted by flow cytometry (BD FACSAria Fusion) on the basis of ATTO 550 fluorescence for clonal culture in 96-well plates (TPP) coated with CellAdhere Laminin-521 (STEMCELL Technologies). After 6-10 days, half of each colony was transferred to Matrigel-coated 24-well plates for expansion, and the remaining cells were lysed in 25 µL of QuickExtract DNA Extraction Solution (Lucigen). Targeted regions were amplified by PCR and Sanger-sequenced to confirm on-target editing (fig.S2A); potential off-target loci predicted to carry two or three mismatches to the gRNA were similarly amplified and sequenced. Four homozygous ($PAX3^{-/-}$, clones 1-4) and three heterozygous ($PAX3^{+/-}$, clones 1-3) mutant clones, all carrying the same single-nucleotide insertion (*c.194_195insA*), were selected for all experiments, together with four WT control lines: the parental WTC-11 line (WT clone 1) and three isogenic WT clones (WT clones 2-4). The resulting frameshift is predicted to yield a truncated PAX3 protein lacking both the paired/homeodomain DNA-binding and C-terminal transactivation domains (fig.S2A). All clones were tested for genomic stability (iCS-digital PSC 24-probe test, Stem Genomics) and for pluripotency by immunostaining for SOX2 and OCT4. All lines tested negative for mycoplasma contamination. More importantly, they were able to differentiate into $SOX1^+$; $PAX6^+$; $PAX7^+$ dorsal spinal progenitors (Fig. 2B and fig. S2C).

Generation of iPSC lines expressing VINCULIN tension sensor (Vin-TS) constructs

*Vin-TS* and $Vin^{T12}$*-TS* construct sequences were PCR-amplified from in-house plasmids described previously (*2*) and cloned into an *iON-pCAG* backbone (*3*) by In-Fusion cloning (Takara Bio) using the XmaI and BamHI restriction sites of an *iON-pCAG-RFP* vector. Primer sequences are listed in Table S1. All constructs were verified by Sanger sequencing. The Escherichia coli MACH1 strain was used for plasmid amplification, and plasmid DNA was prepared using standard miniprep and midiprep kits (Thermo Scientific and Macherey-Nagel). iON plasmids encoding the *Vin-TS* constructs, together with a *piggyBac* expression plasmid (2:1 ratio) (*3*), were transfected into one WT and one $PAX3^{-/-}$ iPSC line using Lipofectamine 3000 (Invitrogen) according to the manufacturer's protocol. Five days after transfection, fluorescent cells were selected by FACS (BD FACSAria Fusion) on the basis of Venus fluorescence after 488-nm excitation and expanded in culture.

Differentiation of iPSCs into spinal organoids

Spinal organoids were generated from iPSCs dissociated into single cells with accutase (STEMCELL Technologies) and seeded in ultra-low-attachment 6-well plates (Corning) in suspension culture, as described previously (*4*, *5*). Cells were cultured in N2B27 differentiation medium (Advanced DMEM/F12 and Neurobasal, 1:1) supplemented with N2, B27 without vitamin A, 1% GlutaMAX, 0.2% β-mercaptoethanol, and 1% penicillin-streptomycin (all from Thermo Fisher Scientific). The ROCK inhibitor Y-27632 (10 µM, Axon Medchem) was included during the first two days to promote cell survival. Medium was refreshed on days 2 and 4. Specification was achieved by modulating signaling pathways with small molecules: Wnt signaling was activated from days 0-4 with CHIR99021 (3-4 µM, Axon Medchem); retinoic acid (10 nM, Sigma) was added on day 2 to promote neural differentiation; and BMP4 (5 ng/mL, R&D Systems) was applied from days 4-7 to induce dorsalization. Small molecules used to modulate glycolysis and the mannose pathway were added to the medium; their concentration, supplier, and treatment window are provided in Table S2. For acute reduction of actomyosin contractility, organoids were treated with the myosin II inhibitor blebbistatin (Sigma); concentration, supplier, and treatment window are provided in Table S2. For glucose-deprivation experiments, organoids were cultured in glucose-free DMEM and glucose- and sodium pyruvate-free Neurobasal-A (both Thermo Fisher Scientific; 1:1), supplemented with

1% sodium pyruvate (Thermo Fisher Scientific), N2, B27 without vitamin A, 1% GlutaMAX, 0.2% β-mercaptoethanol, and 1% penicillin-streptomycin.

Mouse lines

Mice carrying the *Pax3 GFP* knock-in null allele (*Pax3$^{GFP}$*), described previously (*6*), were used. Animals were handled according to European Community guidelines, implementing the 3Rs rules. Protocols were validated by the ethics committee of the French Ministry.

Preparation of micropatterned neuroepithelial culture

*Fabrication of PDMS stamps and PDMS-coated substrates*

Polydimethylsiloxane (PDMS; Sylgard 184, Dow Corning) was mixed with curing agent at a 10:1 ratio, poured onto silicon wafers bearing 700-µm-diameter circular micropatterns produced by photolithography of SU-8 resin, and degassed under vacuum for 30 min. PDMS molds were cured for at least 4 h at 80 °C. Glass coverslips (24 × 60 mm, Brand) or 35-mm culture dishes (ibidi) were coated with PDMS using a spin coater (Laurell) and cured for at least 4 h at 80 °C.

*Microcontact printing of laminin*

CellAdhere Laminin-521 (STEMCELL Technologies) was diluted in PBS containing calcium and magnesium (PBS++; Thermo Fisher Scientific) and applied onto PDMS stamps for at least 1 h under humidified conditions to prevent drying. Stamps were then washed once with PBS++, then with distilled water, and air-dried. PDMS-coated coverslips or dishes were exposed to deep UV for 5 min in a UV-ozone cleaner before contact printing with the laminin-coated stamps. Printed coverslips were mounted in a Well Slip 10-well magnetic chamber (Gataca Systems). 1% pluronic F-127 (Sigma) was then applied for 20 min, followed by three washes with PBS++.

Organoid dissociation and differentiation on micropatterns

Day 4 organoids were dissociated into single cells by incubation with accutase (STEMCELL Technologies) for 10 min at room temperature with regular agitation. Accutase was diluted in Advanced DMEM/F12, and cells were collected by centrifugation and resuspended in N2B27 differentiation medium supplemented with retinoic acid (10 nM, Sigma) and the ROCK inhibitor Y-27632 (10 µM, Axon Medchem). Cells were seeded uniformly onto micropatterned substrates at $5 \times 10^4$ cells per magnetic-chamber well or $2 \times 10^5$ cells per culture dish. After 1 h, the medium was replaced to remove non-adherent cells, and cultures were maintained in N2B27 medium supplemented with retinoic acid (10 nM, Sigma) and recombinant human BMP4 (5 ng/mL, R&D Systems). Where indicated, micropatterned progenitors were treated with mannose and/or the myosin II inhibitor blebbistatin; concentrations, suppliers, and treatment windows are provided in Table S2.

Transmission electron microscopy

Organoids were fixed in 1% glutaraldehyde and 2% paraformaldehyde in PBS (pH 7.4) for 2 h. After washing in 1× PBS, organoids were post-fixed for 1 h in 1% osmium tetroxide reduced with 1.5% potassium ferrocyanide in 1× PBS. Following washes in distilled water, organoids were dehydrated through a graded ethanol series (30%, 50%, 70%, 80%, 90% twice, and 100% three times), each step lasting 10 min. Resin infiltration was carried out with Agar low-viscosity resin (Agar Scientific) mixed with 100% ethanol at ratios of 1:3 for 1 h, 1:2 for 1 h, and 3:4 overnight. The following day, organoids were incubated in pure resin for three successive 2 h baths before embedding in molds. Polymerization was performed overnight at 60 °C. Ultrathin sections (70 nm) were cut with a Leica EM UC6 ultramicrotome, post-stained with 2% aqueous uranyl acetate and 3% lead citrate (Reynolds' solution), and examined at 120 kV on a Tecnai 12 transmission electron microscope equipped with a 4K × 4K Gatan OneView camera.

Immunofluorescence

Mouse embryos were fixed in 4% paraformaldehyde for 1 h at room temperature, cryoprotected by equilibration in 15% sucrose, embedded in gelatin, cryosectioned at 14 µm, and processed for immunostaining as described previously (*7*). Organoid fixation, embedding, and cryosectioning were performed as described previously (*5*), and organoid sections were immunostained using the same protocol as embryo sections. Micropatterned neuroepithelial cultures grown in magnetic chambers were fixed in 4% paraformaldehyde for 30 min at room temperature; the protocol was adapted from that used for organoid sections by adding, after fixation, an overnight incubation at 4 °C in a blocking solution of PBS with 3% bovine serum albumin (Millipore) and 1% Triton X-100 (Sigma). F-actin was labeled with rhodamine phalloidin (1:500, Cytoskeleton) for 10 min on sections and 30 min on micropatterned neuroepithelia. Details of primary and secondary antibodies are provided in Table S3.

Image acquisition and analysis

Fluorescence images were acquired on either a Leica TCS SP5 or a Zeiss LSM 980 confocal microscope; high-resolution imaging used the Airyscan 2 multiplex mode of the LSM 980. The proportion of immunolabeled cells was quantified in CellProfiler (Broad Institute): nuclei were segmented on the basis of DAPI fluorescence using an intensity threshold, and positive cells were expressed as a percentage of the total detected nuclei (e.g. Fig. 1D). For nuclear orientation analyses in Fig. 2G, nuclear segmentation and extraction of geometric features were performed with StarDist in QuPath (*8*, *9*), and nuclear orientation was computed from the second-order geometric moments of the nucleus polygon. Orientation relative to the nearest neighboring nucleus was then computed using a custom Python pipeline, in which a KD-tree was used to efficiently identify the nearest spatial neighbor of each nucleus and to calculate the relative orientation angle between neighboring nuclei. Nuclei within the rosette lumen were excluded from the analysis. Adherens junction components (CDH2, α-catenin, β-catenin) and F-actin distributions along the apico-basal axis were quantified by manually tracing 0.9-µm-wide bands from the apical to the basal surface along cell-cell junctions. For each profile, the cluster extension was defined as the distance required to reach half-maximal fluorescence intensity, and the maximal cluster intensity was recorded; both were extracted with a custom Python pipeline. Apico-basal widths of ZO-1 and PARD3 were measured in the same way. Depending on the figure, cluster extension and intensity are displayed either as per-contact plots with mean ± SEM (e.g. Fig. 3, Fig. 6B) or as polar bar diagrams in which each segment length equals the cluster extension (e.g. fig. S10C). VINCULIN clusters were manually segmented using a graphics tablet, and VINCULIN and phospho-myosin light chain II (pMLCII) fluorescence intensities were quantified in Fiji v2.16.0 (NIH) (e.g. Fig. 4B). Geometrical and conformational properties of two-dimensional neuroepithelia in Fig. 4E and fig. S6F were analyzed with the Tissue Analyzer plugin in Fiji (*10*). Organoid projected area in fig. S2E was measured in Fiji from low-magnification widefield images acquired on a Zeiss Axio Observer microscope. LAMININ fragments were segmented and their geometric properties quantified in Fiji (e.g. Fig. 2E). Scripts used to process the data are available under a GNU General Public License v3.0 on GitHub (*https://github.com/ribeslab/Mirdass-et-al*).

Deep learning-based image classification

Microscopy images were classified into three phenotypic categories (WT, *PAX3*$^{+/-}$, and *PAX3*$^{-/-}$) using a transfer-learning approach based on a ResNet-50 architecture pretrained on the ImageNet dataset (Stanford University) (*11*). Convolutional layers corresponding to residual blocks 1–3 were frozen, whereas the final residual block (layer 4) and the classification head were fine-tuned. The classification head consisted of a dropout layer ($p$ = 0.5) followed by a fully connected layer projecting the 2048-dimensional feature representation onto the three output classes. The dataset was split into training (80%) and validation (20%) sets. Model training was performed using cross-entropy loss and the AdamW optimizer. Data augmentation was applied during training to improve model generalization. Model performance was evaluated by monitoring the loss calculated on the training and validation sets (e.g. fig. S3C) and the checkpoint at epoch 38 was selected for subsequent analysis. Classification accuracy was evaluated by a confusion matrix analysis (e.g. fig. S3D). Latent feature representations extracted from the classifier were analyzed by uniform manifold approximation and projection (UMAP) to visualize the organization of the learned feature space (e.g. Fig. 2D). The scripts used for this study are provided on GitHub (*https://github.com/ribeslab/Mirdass-et-al*).

Laser ablation imaging and quantification

Micropatterned neuroepithelial cultures were generated in ibidi culture dishes and incubated with SPY555-actin (Spirochrome) diluted 1:1000 in FluoroBrite DMEM (Thermo Fisher Scientific) for 2 h to label F-actin. Following incubation, half of the medium was replaced with fresh FluoroBrite DMEM. Laser ablation experiments were performed on a Yokogawa Spinning Disk CSU-X1 microscope maintained at 37°C under 5% $CO_2$. Images were acquired using a 100×/1.4 NA oil-immersion objective. SPY555-actin fluorescence was excited with a 561-nm laser and collected through a 590/35 emission filter using a PRIME 95 sCMOS camera (Teledyne Photometrics). Junctional actin ablation was achieved using a 355-nm pulsed laser controlled through MetaMorph software with a user-defined region of interest encompassing the target junction. Time-lapse imaging was acquired at 100 ms intervals for 30 s following ablation. The displacement of the two tricellular junctions adjacent to the ablation site was quantified by particle image velocimetry using PIVlab. Recoil measurements were subsequently analyzed in Python using a custom script provided on GitHub (*https://github.com/ribeslab/Mirdass-et-al*). Initial recoil velocity was calculated as the distance between the two tricellular junctions at $t_1$ = 0.4s minus the distance at $t_0$ = 0s before the cut, divided by the time between $t_1$ and $t_0$. The recoil velocity curve was calculated after smoothing (using the savgol_filter function on python) and numerically differentiating recoil distance profiles with respect to time.

FRET tension sensors imaging and analysis

Micropatterned neuroepithelial cultures were imaged in a FluoroBrite DMEM medium (Thermo Fisher Scientific) at 37°C and 5% CO2. Histological sections of previously fixed organoids were stained with DAPI (Thermo Fisher Scientific) and imaged at room temperature. FRET-based molecular tension was measured using a sensitized emission protocol (*12*). Briefly, spectral imaging was performed using a confocal microscope (Carl Zeiss LSM 780 or LSM 980) equipped with a 63×/1.4 NA Plan-Apochromat oil immersion objective. The donor of the FRET pair mTFP1/Venus was excited by the 458nm line of an argon laser (for the LSM 780) or with a 445nm diode laser (for the LSM 980); and emission was sampled at a spectral resolution of 9nm within the 462.8-597.1nm range with a Zeiss MBS 458 dichroic (for the LSM 780) or 455.2-614.5nm range with Zeiss MBS 445/561 dichroic (for the LSM 980), on a spectral array of GaAsP detectors.
The FRET index was computed as:

$$I_{FRET} = \frac{I_A}{I_A + I_D}$$

With $I_D$ the background-subtracted intensity in the FRET donor (i.e. mTFP1) channel (489.7-498.7nm for LSM780 and 490.6-499.5nm for LSM980) following donor excitation, and $I_A$ the

background-subtracted intensity in the FRET acceptor (i.e. Venus) channel (525.5-534.5nm for LSM780 and 526.0-534.9nm for LSM980) following donor excitation. The FRET index was calculated in manually-segmented regions of interest encompassing adherens junction clusters. To correct for inter-microscope variability in spectral imaging, the FRET index for each sample was normalised to the mean WT FRET index obtained on the same microscope.
Image analysis and FRET index measurement was performed on Fiji v2.16.0 and Python. Intensity-weighted FRET images were generated in Fiji using a custom script provided on GitHub (*https://github.com/ribeslab/Mirdass-et-al*).

Flow cytometry-based cell cycle analysis

Approximately 10 organoids per condition were dissociated into single cells by incubation with trypsin (Thermo Fisher Scientific) at 37°C for 7-10 min. Enzymatic dissociation was quenched with fetal bovine serum (Thermo Fisher Scientific), and cells were subsequently washed several times with Advanced DMEM/F12 medium. Cells were then resuspended in 250 µL Advanced DMEM/F12, passed through a cell strainer (35µm mesh), and stained with one drop of Hoechst 33342 Ready Flow Reagent (Invitrogen) for 10 min at 37°C. Hoechst 33342 fluorescence was quantified using an Attune CytPix Flow Cytometer (Thermo Fisher Scientific) following laser excitation at 405nm and emission light was collected using a 440/50 band-pass filter. Flow cytometry data were analysed using FlowJo software (version 10.10). Cell cycle phase distribution was determined using the Dean-Jett-Fox model.

Statistical analysis on image analysis

Data are presented as mean ± standard error of the mean (SEM). Statistical analyses were performed in GraphPad Prism using the tests indicated in the figure legends. Sample sizes, statistical tests, and exact *P* values for all graphs are provided in Data S1.

RNA extraction, qPCR and RNA sequencing

Total RNA was extracted with the NucleoSpin RNA kit (Macherey-Nagel) following the manufacturer's instructions, eluted in RNase-free water, and quantified on a DeNovix DS-11 FX-series spectrophotometer. For reverse-transcription quantitative PCR (RT-qPCR), cDNA was synthesized with SuperScript IV reverse transcriptase (Thermo Fisher Scientific) using random primers and oligo(dT). Quantitative PCR was then performed with SYBR Green I Master (Roche) on a Prime Pro 48 real-time PCR system (Techne) or a LightCycler 480 (Roche). qPCR primers were designed with Primer-BLAST (Table S1). Expression levels for each gene at a given time point were measured in biological duplicates or triplicates and normalized to *TBP*. For transcriptomic analysis, RNA was sent to BGI (Hong Kong), where libraries were prepared and their quality assessed on an Agilent 2100 Bioanalyzer. Then 0.2-0.5 µg was sequenced on a DNBSEQ platform (paired-end, 150 bp). After demultiplexing, reads were trimmed and mapped to the GRCh38.p14 human genome (BGI). The RNA-seq data shown in fig. S8E, from day 5 mouse ESC-derived spinal organoids, were published previously (*13*).

Transcriptomic analyses

Pairwise differential expression analysis of the RNA-seq data was performed with DESeq2 (*14*) (BGI). Differentially expressed genes were defined as those with a mean expression > 2 FPKM in at least one group, an absolute fold change > 1.5, and a Benjamini-Hochberg-adjusted q value < 0.01. RNA-seq data from human and mouse spinal organoids are compiled in Data S2. Volcano plots were generated with matplotlib in Python. Enrichment analyses between differentially expressed genes and pathway-representative gene sets were performed using the hypergeometric distribution (phyper function in R). Gene sets were generated on the basis of a literature review or the Kyoto Encyclopedia of Genes and Genomes (KEGG) pathway database and are listed in Data S2. Heatmaps of relative expression were generated with the seaborn.heatmap function in Python. ssGSEA was performed with the R package GSVA, and the corresponding gene sets are listed in Data S2.

Metabolome analysis

After 7 days of differentiation, organoids were washed with cold PBS, snap-frozen in liquid nitrogen, and stored until analysis. A total of 10 biological replicates per condition were generated, corresponding to triplicate or quadruplicate samples from three independent clones per genotype. Samples were subsequently shipped to the Laboratory of Innovations in Mass Spectrometry for Health (Institut des sciences du vivant Frédéric-Joliot, Saclay, France) for metabolomics analyses, as described below.

*Metabolite extraction*

Metabolite extraction was performed on the organoid samples, to which 170 µL of water was added, and an ultrasonic step at 40% amplitude for 10 seconds was applied to each sample. 20 µL of each sample was kept and stored at -20°C for subsequent determination of the total protein concentration (Pierce BCA Protein Assay Kit, Thermo Fisher Scientific). The remaining 150 µL samples were resuspended in 350 µL of methanol containing internal standards at 3.75 µg/mL (dimetridazole, AMPA, MCPA, dinoseb; Sigma). After vortexing, samples were left on ice for 90 minutes until complete protein precipitation. After a final centrifugation step at 20,000 g for 15 minutes at 5°C, supernatants were recovered and split into two equal aliquots of 180 µL for C18 and HILIC analyses. The resulting aliquots were then dried under a stream of nitrogen using a TurboVap instrument (Thermo Fisher Scientific) and stored at -20°C until analysis. Prior to LC-MS analysis, dried extracts were resuspended to reach a fixed protein concentration (equivalent to 5 µg/10 µL) using variable volumes (according to BCA) of H2O/acetonitrile (95:5, v/v) containing 0.1% formic acid + EI* or 10 mM ammonium carbonate pH 10.5 + EI*/acetonitrile (40:60, v/v) for metabolite analysis using C18 and ZIC-pHILIC columns, respectively. After reconstitution, the tubes were vortexed and then centrifuged at 20,000 g for 15 minutes at 5°C. The supernatant was transferred into 0.2 mL vials. A quality control (QC) sample was obtained by pooling about 140 µL of each sample preparation. QC samples were injected every 10 samples in order to evaluate the signal variations of any metabolite. *EI (external standards): mixture of 9 authentic chemical standards covering the mass range of interest (13C-glucose, 15N-aspartate, ethylmalonic acid, amiloride, prednisone, metformin, atropine sulfate, colchicine, imipramine) added to all samples in order to check for consistency of analytical results in terms of signal and retention time stability throughout the experiments

*Liquid chromatography coupled to high-resolution mass spectrometry*

The ultra-high performance liquid chromatographic (UHPLC) separation was performed on a Hypersil GOLD C18 1.9 µm, 2.1 mm x 150 mm column (RP) at 30°C (Thermo Fisher Scientific), and HPLC chromatographic separations were performed on a Sequant ZICpHILIC 5 µm, 2.1 x 150 mm (HILIC) at 15°C (Merck). All chromatographic systems were equipped with an on-line prefilter (Thermo Fisher Scientific). Experimental settings for each LC/MS condition are described below. Mobile phases for RP columns were 100% water in A and 100% ACN in B, both containing 0.1% formic acid. Regarding HILIC, phase A consisted of an aqueous buffer of 10 mM ammonium carbonate in water adjusted to pH 10.5 with ammonium hydroxide, whereas pure ACN was used as solvent B. Chromatographic elutions were achieved under gradient conditions as follows: (i) RP-based system: the flow rate was set at 500 µL/min. The elution consisted of an isocratic step of 2 minutes at 5% phase B, followed by a linear gradient from 5 to 100% of phase B for the next 11 minutes. These proportions were kept constant for 12.5 min before returning to 5% B for 4.5 min. (ii) HILIC-based system: the flow rate was 200 µL/min. Elution started with an isocratic step of 2 min at 80% B, followed by a linear gradient from 80 to 40% of phase B from 2 to 12 min. The chromatographic system was then rinsed for 5 min at 0% B, and the run ended with an equilibration step of 15 min (80% B). LC-MS analyses were performed using a Vanquish liquid chromatography system coupled to an Exploris 240 mass spectrometer from Thermo Fisher Scientific fitted with an electrospray source operated in the positive and negative ion modes. The software interface was Xcalibur (version 4.7) (Thermo Fisher Scientific). The mass spectrometer was calibrated before each analysis in both ESI polarities using the manufacturer's predefined methods and recommended calibration mixture provided by the manufacturer (external calibration). The Exploris 240 mass spectrometer was operated with capillary voltage at -3 kV in the negative ionization mode and 3 kV in the positive ionization mode and a capillary temperature set at 280°C. The sheath gas pressure and the auxiliary gas pressure were set, respectively, at 60 and 20 arbitrary units with nitrogen gas. The mass resolution power of the analyzer was 120,000 m/Δm, full width at half maximum (FWHM) at m/z 200, for singly charged ions. The detection was achieved from m/z 70 to 1000 for RP conditions in the positive ionization mode and from m/z 70 to 1000 for HILIC conditions in the negative ionization mode.

*Data processing*
All raw data were manually inspected using the Qualbrowser module of Xcalibur version 4.7 (Thermo Fisher Scientific). Automatic peak detection and integration were performed using the XCMS (https://github.com/sneumann/xcms) R package using an in-house integrative bioinformatics pipeline workflow, which returned a data matrix containing m/z and retention time values of features together with their concentrations expressed in arbitrary units (i.e., areas of chromatographic peaks). Features were thereafter filtered according to the following criteria: (i) ratio of chromatographic peak area of biological to blank samples above a value of 3, (ii) the correlation between dilution factors of QC samples and areas of chromatographic peaks (filtered variables should exhibit coefficients of correlation above 0.7 in order to account for metabolites occurring at low concentrations and which are not detected anymore in the most diluted samples) and (iii) repeatability (the coefficient of variations obtained for chromatographic peak areas of QC samples should be below 30%).

*Annotation*
Features were annotated by matching their accurate measured masses ± 10 ppm with theoretical masses contained in biochemical and metabolomic databases by using an informatics tool developed in R language. The databases used were the Kyoto Encyclopedia of Genes and Genomes (KEGG) , the Human Metabolome Database (HMDB) and METLIN. Features were additionally annotated against an in-house spectral database made from ~1200 authentic pure standards analyzed under identical conditions. As proposed by the Metabolomics Standards Initiative, to be identified, metabolites had to match at least two orthogonal criteria (accurately measured mass, isotopic pattern, MS/MS spectrum and retention time) to those of the corresponding standard.

*Statistical analyses*
Dimensionality reduction was performed by partial least squares discriminant analysis (PLS-DA) using the ropls package in R. Differential metabolite abundance between experimental groups was assessed using the Mann-Whitney test, with statistical significance defined as p value < 0.05. Heatmaps of relative metabolite abundance were generated using the

seaborn.heatmap function in Python. Hierarchical clustering of samples was performed using the scipy.cluster.hierarchy.linkage function in Python with the Euclidean metric. Metabolomics data are compiled in DataS3.

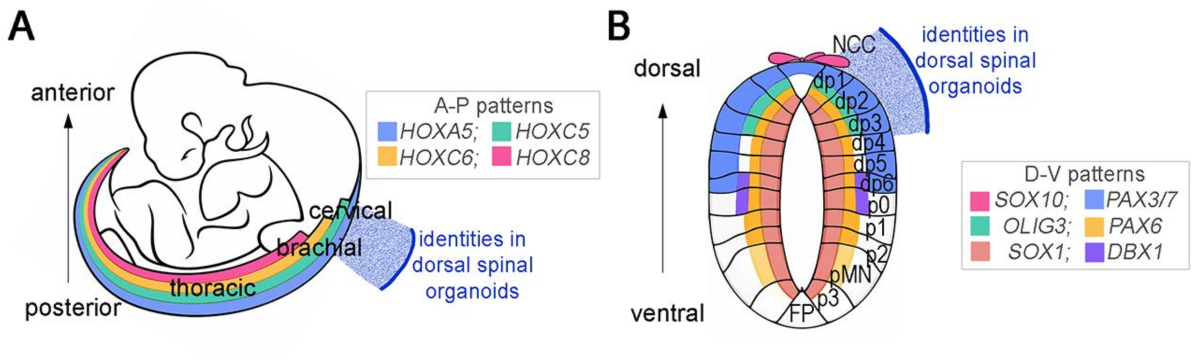


**Fig. S1. Anteroposterior and dorsoventral positional identities represented in dorsal spinal cord organoids.**

**(A)** Schematic of a Carnegie stage 14 human embryo illustrating anteroposterior (AP) patterning of the developing spinal cord and the corresponding *HOX* gene expression domains. **(B)** Schematic cross-section of the neural tube illustrating dorsoventral (DV) patterning and the major progenitor domains of the developing spinal cord. Positional identities represented in dorsal spinal cord organoids are indicated by a blue arc in both panels.

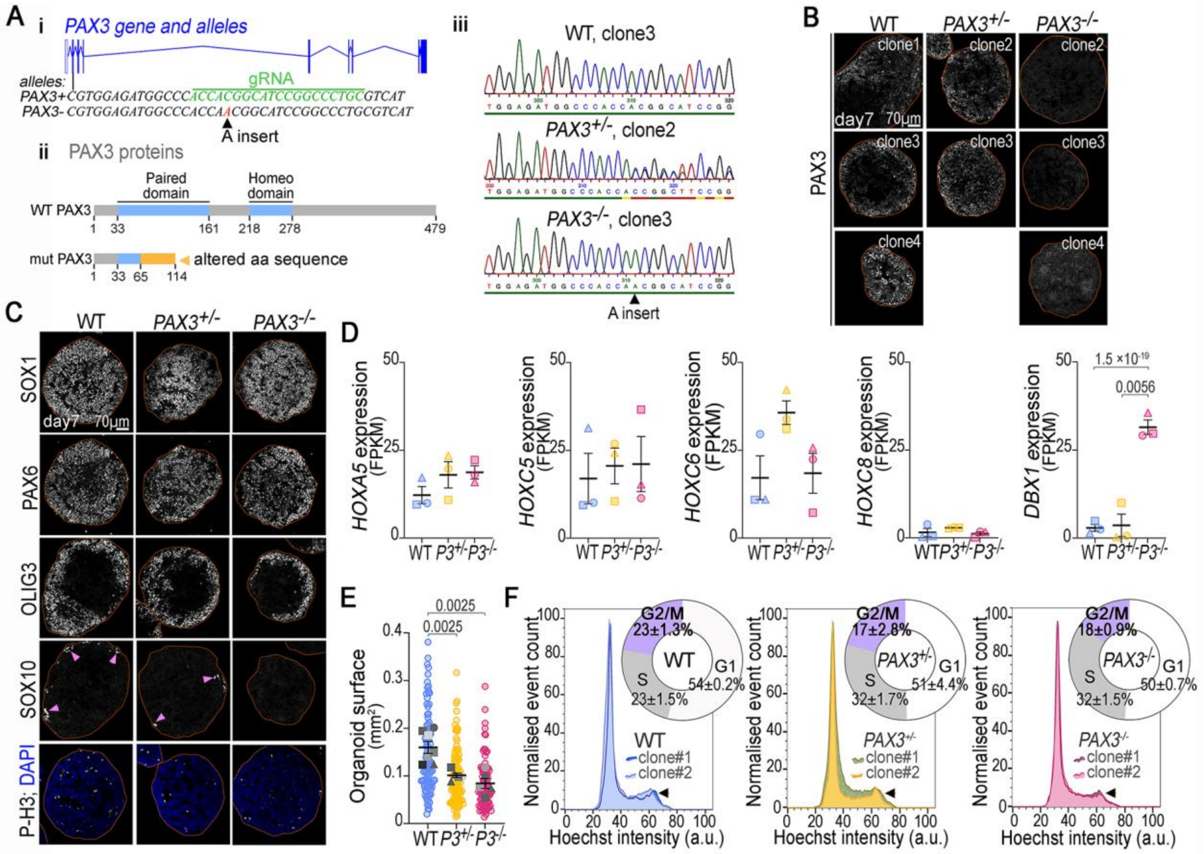


**Fig. S2. Neural identity and growth properties of dorsal spinal cord organoids upon loss of one or both *PAX3* alleles.**

**(A)** Schematic of the WT *PAX3* allele and of the CRISPR-engineered mutant allele carrying a single adenine insertion in exon 2 (*c.194_195insA*) (i); predicted WT and mutant PAX3 protein structures (ii); and sequences of PCR-amplified genomic regions flanking the insertion site from representative WT, *PAX3*$^{+/-}$, and *PAX3*$^{-/-}$ iPSC clones used in this study (iii). **(B)** Representative images of day 7 organoids from independent WT, *PAX3*$^{+/-}$, and *PAX3*$^{-/-}$ iPSC clones (distinct from those in Fig. 2A) stained for PAX3. Orange dashed lines, organoid boundaries. **(C)** Representative images of day 7 WT, *PAX3*$^{+/-}$, and *PAX3*$^{-/-}$ organoids stained for the neural progenitor markers SOX1 and PAX6, the dorsal interneuron progenitor marker OLIG3, the neural crest cell marker SOX10, and the mitotic marker phospho-histone H3 (PH3). Orange dashed lines, organoid boundaries. **(D)** *HOX* gene and *DBX1* expression in day 7 WT, *PAX3*$^{+/-}$, and *PAX3*$^{-/-}$ organoids, assessed by RNA-seq and expressed as fragments per kilobase of transcript per million mapped reads (FPKM). Dots, individual clones; bars, mean ± SEM. DESeq2 adjusted *P* values are shown (statistical details in Data S2). **(E)** Projected area of day 7 organoids measured from low-magnification bright-field images. Colored dots, individual organoids; gray dots, per-clone means within each experiment (symbol shape indicates clone); bars, mean ± SEM across independent clone/experiment replicates. Mann-Whitney *P* values are shown; individual data points, sample sizes, and statistical analyses are provided in Data S1 **(F)** Flow cytometry plots of DNA content distributions of Hoechst 33342-stained cells from two independent clones per genotype, and circular diagrams showing percentage of cells in the indicated cell cycle phases in day 7 WT, *PAX3*$^{+/-}$, and *PAX3*$^{-/-}$ organoids; values, mean ± SEM. Individual values in Data S1.

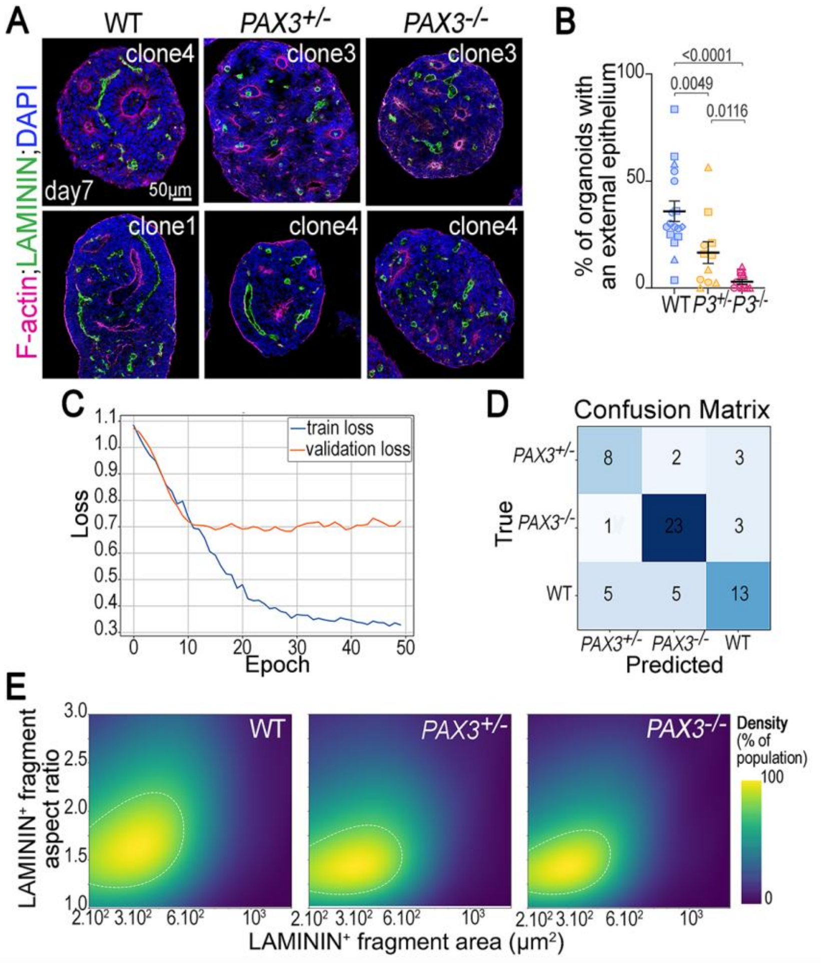


**Fig. S3. Quantitative characterization of neuroepithelial abnormalities in *PAX3*-mutant dorsal spinal cord organoids.**

**(A)** Representative images of day 7 WT, *PAX3*$^{+/-}$, and *PAX3*$^{-/-}$ organoids (clones distinct from those in Fig. 2C) stained for LAMININ, F-actin (phalloidin), and DAPI. **(B)** Percentage of day 7 organoids displaying peripheral epithelia in the indicated genotypes. Dots, individual clone/experiment replicates (symbol shape indicates clone); bars, mean ± SEM. Mann-Whitney *P* values are shown; individual data points, sample sizes, and statistical analyses are provided in Data S1. **(C)** Training and validation loss curves for the ResNet-50 classifier trained on images of day 7 organoids stained for LAMININ, F-actin, and DAPI. **(D)** Confusion matrix showing the classification performance of the ResNet-50 model on previously unseen images of day 7 WT ($n$ = 23), *PAX3*$^{+/-}$ ($n$ = 13), and *PAX3*$^{-/-}$ ($n$ = 27) organoids. **(E)** Kernel density estimates of LAMININ fragment aspect ratio as a function of fragment area in day 7 WT, *PAX3*$^{+/-}$, and *PAX3*$^{-/-}$ organoids, shown for a representative clone of each genotype. The color scale indicates the probability density of fragment occurrence, from dark blue (low density) to yellow (high density); estimates for all clones are provided in Data S1. Dashed contours correspond to 70% of the maximum kernel density estimate and match those shown in Fig. 2E.

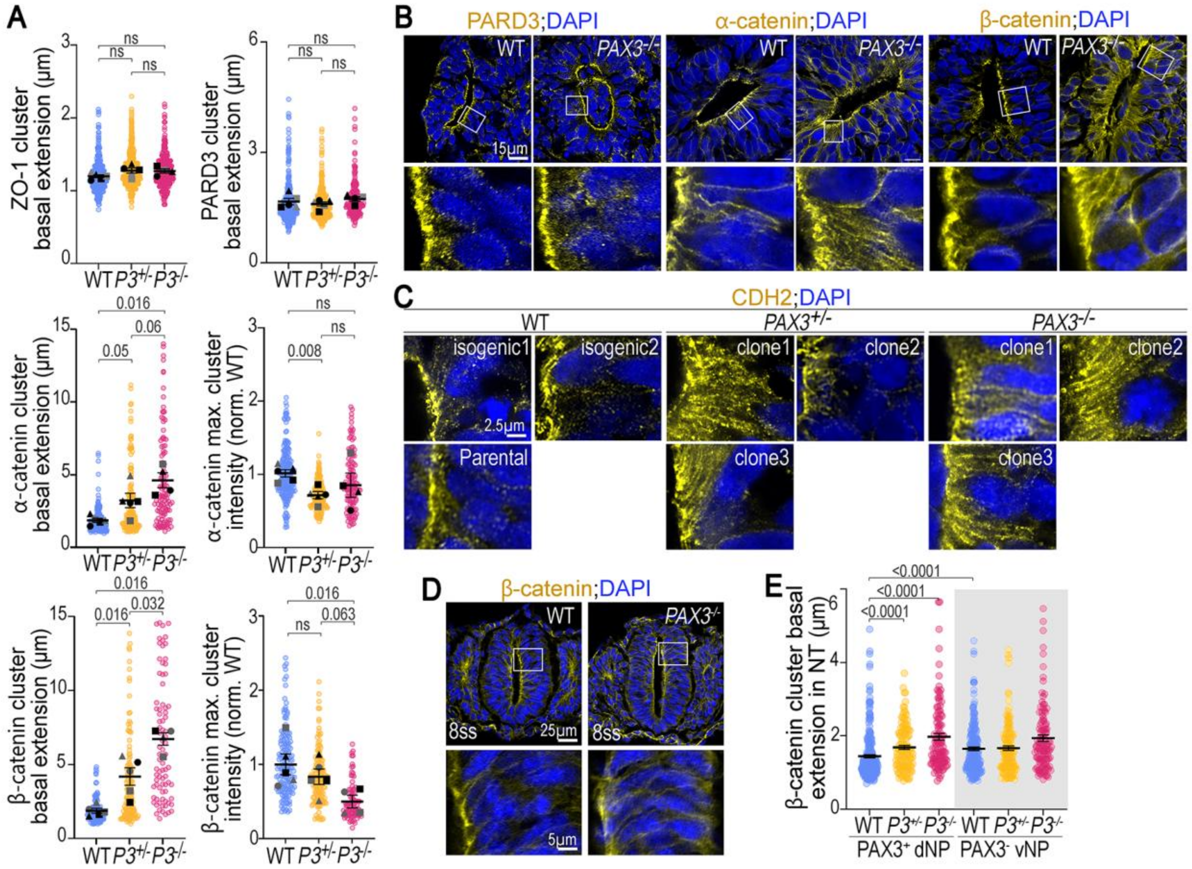


**Fig. S4. Basal expansion of adherens junction domains and preserved apical polarity in *PAX3*-mutant neural epithelia.**

**(A)** Apical marker distributions in day 7 WT, *PAX3*$^{+/-}$, and *PAX3*$^{-/-}$ organoid rosettes. Cluster extension along the apical-basal axis for ZO-1 (top left), PARD3 (top right), α-catenin (middle left), and β-catenin (bottom left); maximum cluster intensity normalized to WT for α-catenin (middle right) and β-catenin (bottom right). Colored dots, individual clusters; gray dots, per-experiment means (symbol shape indicates clone); bars, mean ± SEM across independent clone/experiment replicates. **(B)** Representative images of day 7 WT, *PAX3*$^{+/-}$, and *PAX3*$^{-/-}$ organoid rosettes stained for PARD3, α-catenin, β-catenin, and DAPI; white boxes, regions shown at higher magnification below. **(C)** Representative higher-magnification images of apical domains from day 7 WT, *PAX3*$^{+/-}$, and *PAX3*$^{-/-}$ organoid rosettes, across independent clones, stained for CDH2 and DAPI. **(D)** Representative images of E8.5 WT and *Pax3*$^{GFP/GFP}$ mouse embryos at the level of somite 5, stained for β-catenin and DAPI; white boxes, regions shown at higher magnification below. **(E)** Apical β-catenin cluster extension along the apical-basal axis in mouse embryos of the indicated genotypes, measured in the dorsal half (white background) or ventral half (gray background) of the neural tube. Dots, individual clusters ($n$ = 4 embryos per genotype); bars, mean ± SEM across independent clusters. For all quantifications, Mann-Whitney $P$ values are shown on the plots; individual data points, sample sizes, and statistical analyses are provided in Data S1.

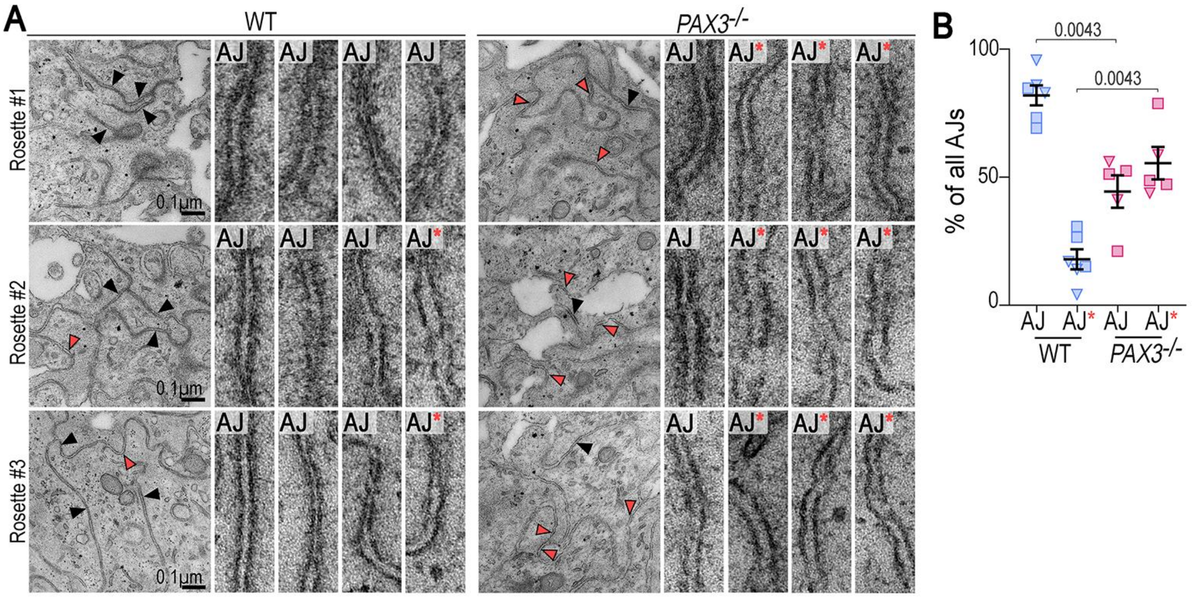


**Fig. S5. Ultrastructural adherens junction defects in *PAX3*-mutant dorsal spinal cord organoids.**

**(A)** Representative TEM images of three-day 7 WT and *PAX3*$^{-/-}$ organoid rosettes. Left, lower-magnification overviews in which arrows mark individual adherens junctions (AJs); right, higher-magnification views of the four arrowed AJs. Arrows are color-coded: black, normal AJs; orange outlined in black, aberrant AJs (AJ*). **(B)** Percentage of normal (AJ) and aberrant (AJ*) morphologies among all AJs identified in individual day 7 WT and *PAX3*$^{-/-}$ organoid rosettes. Dots, individual rosettes (symbol shape indicates organoid); bars, mean ± SEM across independent rosettes. Mann-Whitney *P* values are shown on the plots; individual data points, sample sizes, and statistical analyses are provided in Data S1.

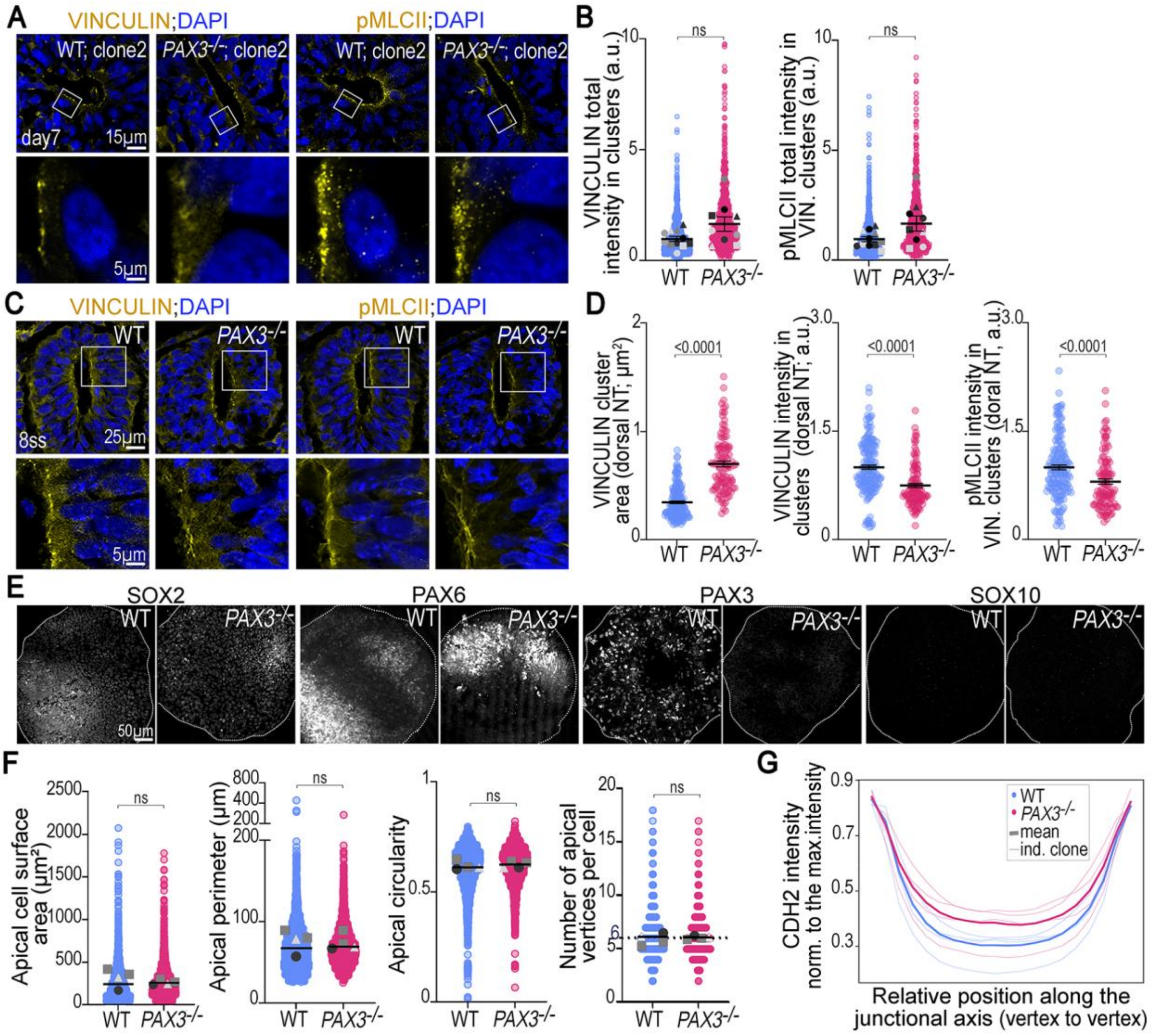


**Fig. S6. Apical junctional disorganization in *PAX3*-mutant organoid, embryonic, and two-dimensional neural epithelia.**

**(A)** Representative images of day 7 WT and $PAX3^{-/-}$ organoid rosettes (clones distinct from those in Fig. 4A) stained for VINCULIN, phosphorylated myosin light chain II (pMLCII), and DAPI; white boxes, regions shown at higher magnification below. **(B)** Total VINCULIN and pMLCII fluorescence intensity within VINCULIN-positive clusters. Colored dots, individual clusters; gray dots, per-experiment means (symbol shape indicates clone); bars, mean ± SEM across independent clone/experiment replicates. **(C)** Representative images of E8.5 WT and *Pax3*$^{GFP/GFP}$ embryos at somite level 5, stained as in (A); white boxes, apical surface of the dorsal neural tube shown at higher magnification below. **(D)** Apical VINCULIN and pMLCII signals in E8.5 dorsal neural tubes. Left to right: VINCULIN-positive cluster area; mean VINCULIN intensity within clusters; pMLCII intensity within VINCULIN-positive clusters. Dots, individual clusters ($n$ = 4 embryos per genotype); bars, mean ± SEM. **(E)** Representative images of day 7 WT and $PAX3^{-/-}$ neural progenitors cultured on micropatterns and stained for SOX2, PAX6, PAX3, and SOX10. **(F)** Apical cell geometry in the honeycomb-like CDH2 junctional network of day 7 WT and $PAX3^{-/-}$ neural progenitors cultured on micropatterns. Left to right: apical surface area, perimeter, circularity, and number of vertices per cell. Colored dots, individual cells; gray dots, per-experiment means; bars, mean ± SEM across independent clone/experiment replicates. **(G)** CDH2 fluorescence intensity profiles along the junctional axis between adjacent tricellular vertices in day 7 neural progenitors cultured on micropatterns. Thick lines, mean across all measurements; thin lines, per-clone replicates. Mann-Whitney *P* values are shown on the plots; individual data points, sample sizes, and statistical analyses are provided in Data S1.

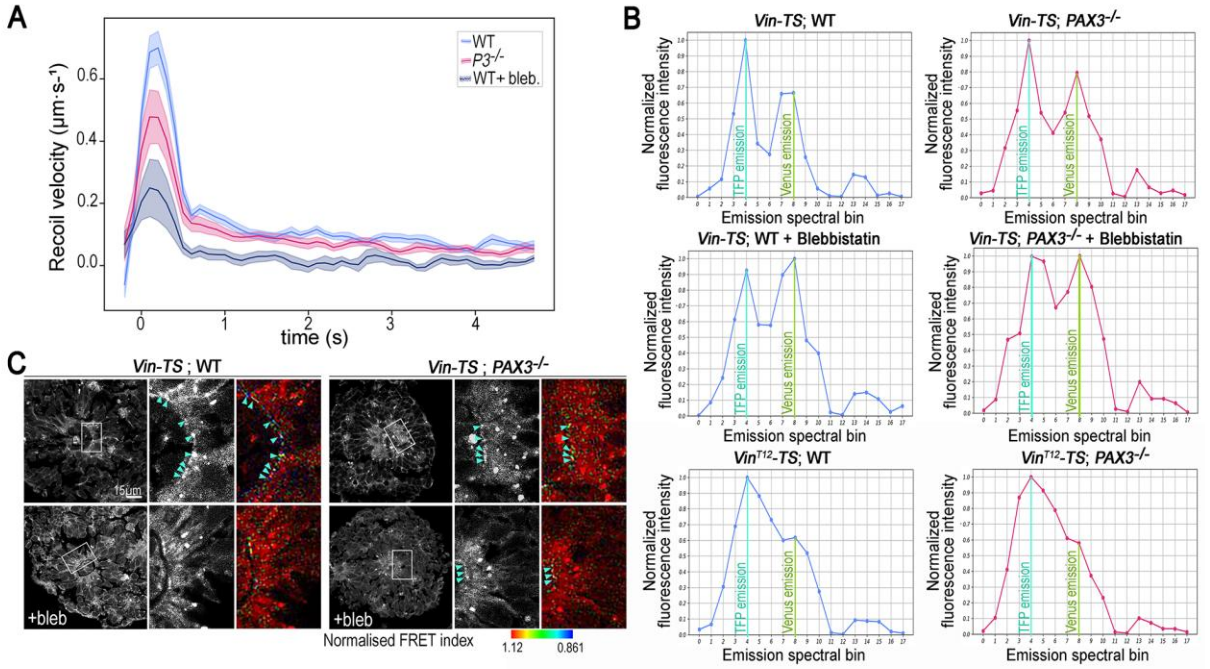


**Fig. S7. Junctional tension measured by laser ablation and validation of the VINCULIN tension sensor (Vin-TS) under blebbistatin-induced tension reduction.**

**(A)** Recoil dynamics following laser ablation of a cell-cell junction labelled with SiR-actin, in WT, blebbistatin-treated WT (WT +Bleb) and *PAX3*$^{-/-}$ neural progenitors at day 7 of differentiation, cultured on micropatterns. Rather than the recoil itself, its smoothed time derivative (µm·s$^{-1}$) is plotted against time after ablation, so that the initial recoil velocity, the readout of relative junctional tension, appears as a distinct peak. Thick lines, mean; thin lines, SEM; individual data points, sample sizes, and statistical analyses are provided in Data S1. **(B)** Fluorescence emission spectra, normalized to maximal intensity, as a function of emission wavelength in WT and *PAX3*$^{-/-}$ neural progenitors cultured on micropatterns and expressing either Vin-TS or Vin$^{T12}$-TS, treated with blebbistatin for 10 min (+Bleb) or left untreated. The donor (TFP) and acceptor (Venus) emission peaks are highlighted in turquoise and green, respectively. **(C)** Representative images of fixed day 7 WT and *PAX3*$^{-/-}$ organoid rosettes expressing Vin-TS, treated with blebbistatin for 10 min (+Bleb) or left untreated. For each genotype and treatment, three panels are shown: a grey low-magnification overview, in which a white box marks the region magnified at right; and two higher-magnification views of that region, one in grey showing Vin-TS signal intensity and one color-coded for normalized FRET index (color scale at right). Turquoise arrowheads, Vin-TS-positive apical clusters used for FRET-index quantification.

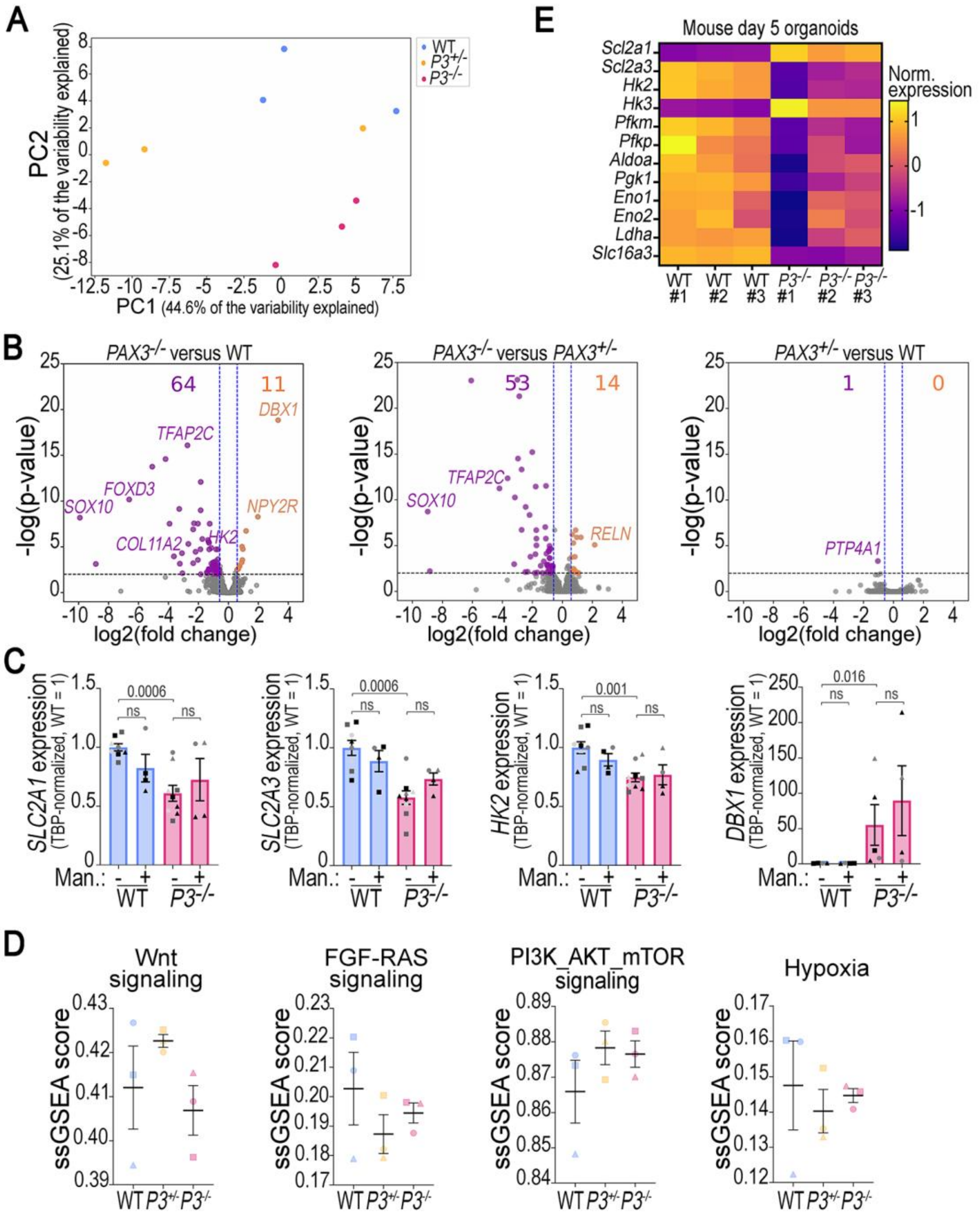


**Fig. S8. Transcriptomic alterations in *PAX3*-mutant spinal cord progenitors.**
**(A)** Principal component analysis (PCA) of gene expression profiles obtained by RNA-seq (FPKM values) from day 7 WT, *PAX3*$^{+/-}$, and *PAX3*$^{-/-}$ organoids. **(B)** Volcano plots of pairwise differential gene expression analyses between *PAX3*$^{-/-}$ and WT organoids (left), *PAX3*$^{-/-}$ and *PAX3*$^{+/-}$ organoids (middle), and *PAX3*$^{+/-}$ and WT organoids (right). Differentially expressed genes are highlighted in orange (upregulated) and purple (downregulated). **(C)** qRT-PCR validation of altered expression of glycolysis-associated genes and *DBX1* in day 7 organoids independent of those analyzed by RNA-seq, treated with or without mannose. Dots, individual experiments (symbol shape indicates clone); bars, mean ± SEM. **(D)** Single-sample gene set enrichment analysis (ssGSEA) scores for gene sets associated with Wnt signaling, FGF-RAS signaling, PI3K-AKT-mTOR signaling, and hypoxia in day 7 WT, *PAX3*$^{+/-}$, and *PAX3*$^{-/-}$ RNA-seq samples. **(E)** Heatmap of the expression of glycolysis-associated genes in day 5 WT and *PAX3*$^{-/-}$ mouse dorsal spinal cord organoids analyzed by

RNA-seq (*n* = 3 biological replicates per genotype) and previously published in (*13*). For (A), (B), and (D), *n* = 3 clones per genotype; sample descriptions, sizes, and ssGSEA signatures are provided in Data S2. Mann-Whitney P values are shown on the (D); individual data points, sample sizes, and statistical analyses in (D) and (E) are provided in Data S1.

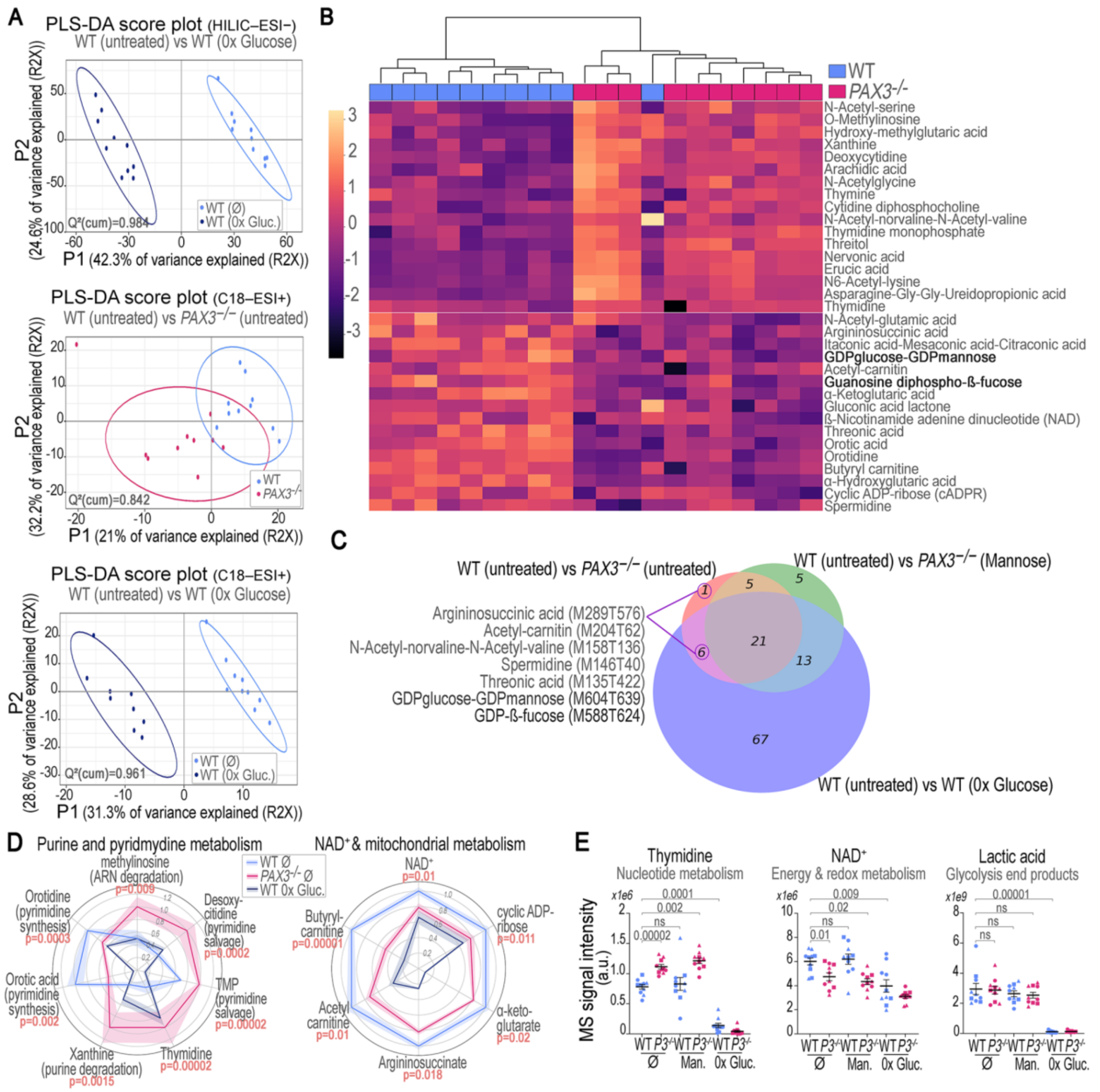


**Fig. S9. Metabolic remodeling induced by *PAX3* loss, glucose deprivation, and mannose supplementation.**

**(A)** PLS–DA score plots of HILIC–ESI– and C18–ESI+ metabolomic profiles from day 7 WT and *PAX3*$^{-/-}$ organoids. $Q^2$(cum) values are indicated for each analysis. **(B)** Heatmaps of metabolites differentially abundant between day 7 WT and *PAX3*$^{-/-}$ organoids, with hierarchical clustering of individual samples ($n$ = 10 samples per genotype and 3 clones per genotype). **(C)** Venn diagram of the overlap between metabolites differentially abundant in the indicated comparisons: WT versus *PAX3*$^{-/-}$ organoids under standard culture conditions; WT versus *PAX3*$^{-/-}$ organoids cultured with mannose supplementation; and WT organoids cultured in standard versus glucose-free medium (0× glucose). The seven metabolites whose abundance was restored in *PAX3*$^{-/-}$ organoids after mannose supplementation are highlighted. **(D)** Polar bar diagrams of the relative mass-spectrometric intensity of the indicated metabolites in day 7 WT (blue) and *PAX3*$^{-/-}$ (pink) organoids, and in WT organoids cultured in glucose-free medium (dark blue). Each radial segment corresponds to one metabolite. For each metabolite, intensities are normalized to the maximum across the three conditions (maximum = 1). **(E)** Relative mass-spectrometric intensity of the indicated metabolites in day 7 WT and *PAX3*$^{-/-}$ organoids, with or without mannose supplementation, and in glucose-free culture. Dots, individual samples; bars, mean ± SEM across independent samples. For (D) and (E), Mann-Whitney *P* values are shown; sample sizes and statistical analyses are provided in Data S3.

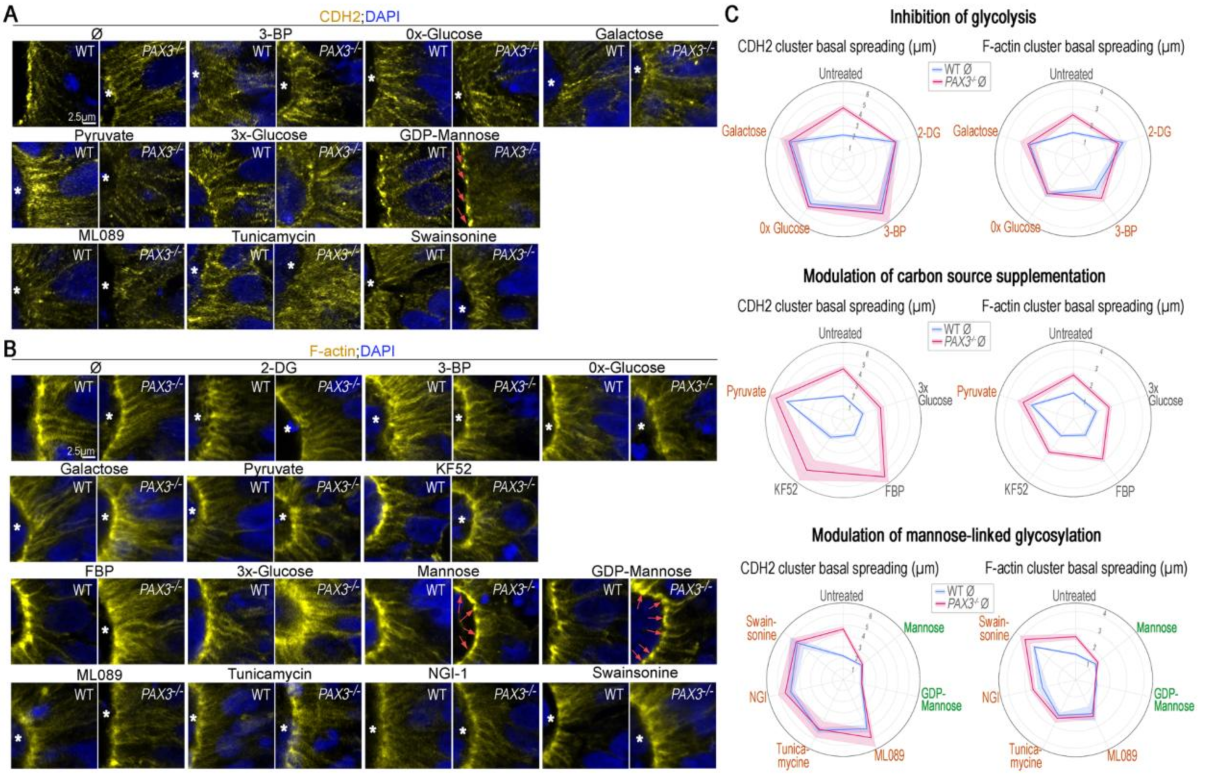


**Fig. S10. Glycolysis- and mannose metabolism-dependent regulation of adherens junction organization in the presence or absence of *PAX3*.**

**(A and B)** Representative high-magnification images of the apical surface of day 7 WT and *PAX3*$^{-/-}$ organoid rosettes treated with the indicated modulators of glycolysis and mannose metabolism and stained for CDH2 (A) or F-actin (phalloidin) (B), and DAPI. Asterisks, regions showing reduced apical CDH2 and F-actin clustering; pink arrows, restoration of apical CDH2 and F-actin clustering in *PAX3*$^{-/-}$ organoids after mannose or GDP-mannose treatment. **(C)** Polar bar diagrams of apical CDH2 (left) and F-actin (right) cluster extension along the apical-basal axis in day 7 WT (blue) and *PAX3*$^{-/-}$ (pink) organoids treated with the indicated modulators of glycolysis and mannose metabolism. Each radial segment corresponds to one treatment, with segment length equal to cluster extension (µm). Treatment labels are color-coded by effect: green, significant rescue of the mutant phenotype; ochre, significant induction of a mutant-like phenotype in WT organoids; gray, no significant change relative to untreated. Individual data points, sample sizes, and statistical analyses are provided in Data S1.

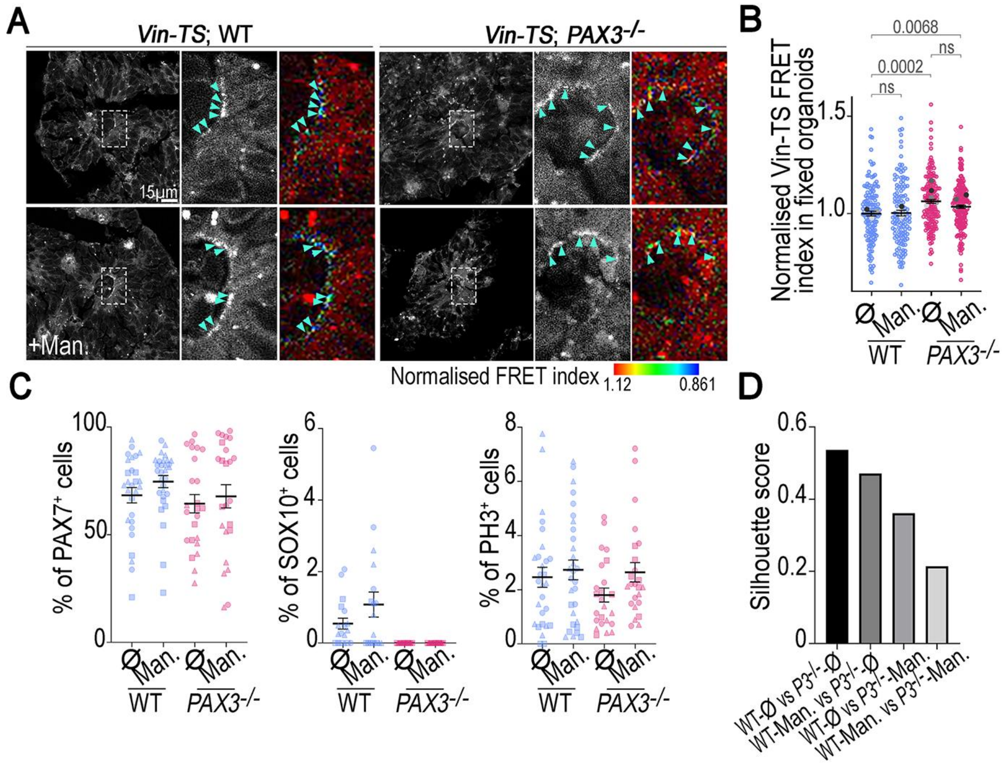


**Fig. S11. Mannose-mediated rescue of *PAX3* loss-induced defects in VINCULIN tension and proliferation, without restoration of cell fate specification.**

**(A)** Representative images of fixed day 7 WT and *PAX3*$^{-/-}$ organoid rosettes expressing Vin-TS, treated with or without mannose (+Man). For each genotype and treatment, three panels are shown: a grey low-magnification overview, in which a dashed box marks the region magnified at right; and two higher-magnification views of that region, one in grey showing Vin-TS signal intensity and one color-coded for normalized FRET index (color scale at right). Turquoise arrowheads, Vin-TS-positive apical clusters used for FRET-index quantification. **(B)** FRET index, normalized to WT (WT = 1), within apical VINCULIN-positive clusters in samples as in (A). Colored dots, individual VINCULIN-positive clusters; gray dots, per-experiment means; bars, mean ± SEM across experiments. **(C)** Percentage of PAX7$^+$, SOX10$^+$, and phospho-histone H3$^+$ (PH3$^+$) nuclei in day 7 WT and *PAX3*$^{-/-}$ organoids treated with or without mannose. Dots, individual organoids (symbol shape indicates clone); bars, mean ± SEM across independent organoids. **(D)** Silhouette scores calculated between pairs of clusters from the UMAP shown in Fig. 6G. Mann-Whitney *P* values are shown on the plots; individual data points, sample sizes, and statistical analyses are provided in Data S1.

**Table S1. Primer sequences.**
Columns indicate the application, primer name, and sequence (5′→3′). DNA sequences are shown in italics.

| Application | Primer name | Sequence (5′→3′) |
|---|---|---|
| *PAX3* genotyping | *PAX3* forward | *TTGCCCCATTTGCTGTCTTT* |
| | *PAX3* reverse | *GCCTTTACGCACCTTCACAA* |
| CRISPR off-target validation | 2-mismatch gRNA off-target *PAX7* forward | *GGTCTTCATCAATGGCGAC* |
| | 2-mismatch gRNA off-target *PAX7* reverse | *CTGGTGCCAACGAGGTAGTAAG* |
| | 2-mismatch gRNA off-target *ADRA2C* forward | *CGCGGGCAGGTTTGAC* |
| | 2-mismatch gRNA off-target *ADRA2C* reverse | *AACGAGAAGGGCATGACCAG* |
| *Vin-TS* cloning (iON vector) | *IonCag-XmaI-VinculinTS* forward | *AAGTGATTAGCCCGGATGCCCGTCTTCCACACG* |
| | *IonCag-BamHI-VinculinTS* reverse | *CACTAAAGTCGGATCTTACTGATACCATGGGGTCTTTCTG* |
| qRT-PCR | *TBP* forward | *CACGAACCACGGCACTGATT* |
| | *TBP* reverse | *TTTTCTTGCTGCCAGTCTGGAC* |
| | *SLC2A1* forward | *GCTCCCTGCAGTTTGGCTAC* |
| | *SLC2A1* reverse | *ATAGCGGTGGACCCATGTCT* |
| | *SLC2A3* forward | *ACTTGAATTAGATTACAGCGATGGG* |
| | *SLC2A3* reverse | *AGCCAAATTGGAAAGAGCCGA* |
| | *HK2* forward | *CACGGAGCTCAACCATGACC* |
| | *HK2* reverse | *CCTTGCGGAACCGCTTAGAG* |
| | *GPI* forward | *CCCGAGTCCTCCCTGTTCAT* |
| | *GPI* reverse | *GAGAAACCACTCCTTCGCCG* |
| | *DBX1* forward | *CCGGCTGGTAGGTGTCTCT* |
| | *DBX1* reverse | *GGCCACGATTCCCCACAAAG* |

**Table S2. Small molecules used for organoid culture and metabolic modulation.** Columns indicate the reagent name, supplier, working concentration, and treatment window, where applicable.

| Reagent | Supplier (catalog no.) | Working concentration | Treatment window |
|---|---|---|---|
| 2-Deoxy-D-glucose (2-DG) | Sigma (D8375) | 5 mM | Days 5-7 |
| 3-Bromopyruvate (3-BP) | MedChemExpress (HY-19992) | 50 µM | Days 5-7 |
| Blebbistatin | Sigma (203390) | 50 µM | 10 mn at day 7 |
| BMP4 | R&D Systems | 5 ng/mL | Days 4-7 |
| CHIR99021 | Axon Medchem | 3-4 µM | Days 0-4 |
| D-Glucose | Sigma (A16828) | 3X (60 mM) | Days 2-7 |
| D-Mannose | Sigma (M2069-25G) | 10 mM | Days 2-7 |
| Fructose-1,6-bisphosphate (FBP) | Santa Cruz Biotechnology (CAS 488-69-7) | 10 mM | Days 6-7 |
| Galactose | Sigma (A12813) | 20 mM | Days 2-7 |
| GDP-mannose | MedChemExpress (HY-N7389A) | 100 µM | Days 2-7 |
| KF52 | MedChemExpress (HY-170404) | 5 µM | Days 5-7 |
| ML089 | MedChemExpress (HY-138802) | 30 µM | Days 5-7 |
| NGI-1 | MedChemExpress (HY-117383) | 10 µM | Days 6-7 |
| Pyruvate | Gibco (11360070) | 3X (1.8 mM) | Days 2-7 |
| Retinoic acid | Sigma | 10 nM | Day 2 |
| Swainsonine (SWA) | Sigma (S8195) | 75 µM | Days 2-7 |
| Tunicamycin | MedChemExpress (HY-A0098) | 1 µM | Days 6-7 |
| Y-27632 (dihydrochloride) | Axon Medchem (#1683) | 10 µM | Days 0-2 |

**Table S3. Antibodies used in this study.**
Columns indicate the target antigen, host species, dilution, supplier (catalogue number), and Research Resource Identifier (RRID).

| Target antigen | Host species | Dilution | Supplier (catalogue no.) | RRID |
|---|---|---|---|---|
| **Primary antibodies** | | | | |
| α-Catenin | Mouse | 1:200 | Enzo Life Sciences (ALX-804-101) | AB_2230168 |
| β-Catenin (CTNNB1) | Rabbit | 1:100 | Abcam (ab16051) | AB_443301 |
| Laminin | Rabbit | 1:500 | Sigma (L9393) | AB_477163 |
| N-Cadherin (CDH2) | Mouse | 1:200 | Sigma (C2542) | AB_258801 |
| OCT3/4 | Mouse | 1:100 | Santa Cruz Biotechnology (sc5279) | AB_628051 |
| OLIG3 | Guinea pig | 1:10,000 | Gift from T. Müller / C. Birchmeier | AB_2315006 |
| PARD3 | Rabbit | 1:500 | Millipore Sigma (07-330) | AB_2101325 |
| PAX3 | Mouse | 1:10 | DSHB (Pax3) | AB_528426 |
| PAX6 | Mouse | 1:50 | DSHB (Pax6) | AB_528427 |
| PAX7 | Mouse | 1:10 | DSHB (Pax7) | AB_528428 |
| Phospho-Histone H3 (Ser10) | Rabbit | 1:500 | Millipore Sigma (06-570) | AB_310177 |
| Phospho-Myosin light chain 2 (Ser19) | Rabbit | 1:100 | Cell Signaling (3671) | AB_330248 |
| SOX1 | Goat | 1:200 | R&D Systems (AF3369) | AB_2239879 |
| SOX2 | Goat | 1:500 | R&D Systems (AF2018) | AB_355110 |
| SOX2 | Mouse | 1:200 | Thermo Fisher Scientific (48-1400) | AB_2533841 |
| SOX10 | Rabbit | 1:2000 | Cell Signaling (89356) | AB_2792980 |
| VINCULIN | Mouse | 1:500 | Millipore Sigma (V9131) | AB_477629 |
| ZO-1 | Rabbit | 1:1000 | Abcam (ab216880) | AB_2909434 |
| **Secondary antibodies** | | | | |
| Goat IgG | Donkey | 1:500 | Thermo Fisher Scientific (A-11056) | AB_2534103 |
| Guinea pig IgG | Donkey | 1:500 | Jackson ImmunoResearch (706-546-148) | AB_2340473 |
| Mouse IgG | Donkey | 1:500 | Thermo Fisher Scientific (A-21202) | AB_141607 |
| Mouse IgG | Donkey | 1:500 | Thermo Fisher Scientific (A10037) | AB_11180865 |
| Mouse IgG | Donkey | 1:500 | Thermo Fisher Scientific (A32787) | AB_2762830 |
| Rabbit IgG | Donkey | 1:500 | Thermo Fisher Scientific (A10042) | AB_2534017 |
| Rabbit IgG | Donkey | 1:500 | Thermo Fisher Scientific (A31573) | AB_2536183 |

**References:**


1. J.-P. Concordet, M. Haeussler, CRISPOR: intuitive guide selection for CRISPR/Cas9 genome editing experiments and screens. *Nucleic Acids Res.* **46**, W242–W245 (2018).

2. H. Canever, H. Lachuer, Q. Delaunay, F. Sipieter, N. Audugé, P. P. Girard, N. Borghi, Collective directional memory controls the range of epithelial cell migration. *eLife* **15** (2026).

3. T. Kumamoto, F. Maurinot, R. Barry-Martinet, C. Vaslin, S. Vandormael-Pournin, M. Le, M. Lerat, D. Niculescu, M. Cohen-Tannoudji, A. Rebsam, K. Loulier, S. Nedelec, S. Tozer, J. Livet, Direct Readout of Neural Stem Cell Transgenesis with an Integration-Coupled Gene Expression Switch. *Neuron* **107**, 617-630.e6 (2020).

4. N. Duval, C. Vaslin, T. Barata, Y. Frarma, V. Contremoulins, X. Baudin, S. Nédélec, V. Ribes, BMP4 patterns Smad activity and generates stereotyped cell fate organisation in spinal organoids. *Development*, dev.175430 (2019).

5. Y. Maury, J. Côme, R. A. Piskorowski, N. Salah-Mohellibi, V. Chevaleyre, M. Peschanski, C. Martinat, S. Nedelec, Combinatorial analysis of developmental cues efficiently converts human pluripotent stem cells into multiple neuronal subtypes. *Nat. Biotechnol.* **33**, 89–96 (2015).

6. A. Zalc, R. Rattenbach, F. Auradé, B. Cadot, F. Relaix, Pax3 and Pax7 Play Essential Safeguard Functions against Environmental Stress-Induced Birth Defects. *Dev. Cell* **33**, 56–66 (2015).

7. J. Briscoe, A. Pierani, T. M. Jessell, J. Ericson, A Homeodomain Protein Code Specifies Progenitor Cell Identity and Neuronal Fate in the Ventral Neural Tube. *Cell* **101**, 435–445 (2000).

8. P. Bankhead, M. B. Loughrey, J. A. Fernández, Y. Dombrowski, D. G. McArt, P. D. Dunne, S. McQuaid, R. T. Gray, L. J. Murray, H. G. Coleman, J. A. James, M. Salto-Tellez, P. W. Hamilton, QuPath: Open source software for digital pathology image analysis. *Sci. Rep.* **7**, 16878 (2017).

9. U. Schmidt, M. Weigert, C. Broaddus, G. Myers, "Cell Detection with Star-Convex Polygons" in *Medical Image Computing and Computer Assisted Intervention – MICCAI 2018: 21st International Conference, Granada, Spain, September 16-20, 2018, Proceedings, Part II* (Springer-Verlag, Berlin, Heidelberg, 2018; https://doi.org/10.1007/978-3-030-00934-2_30), pp. 265–273.

10. B. Aigouy, D. Umetsu, S. Eaton, "Segmentation and Quantitative Analysis of Epithelial Tissues" in *Drosophila*, C. Dahmann, Ed. (Springer New York, New York, NY, 2016; http://link.springer.com/10.1007/978-1-4939-6371-3_13)vol. 1478 of *Methods in Molecular Biology*, pp. 227–239.

11. A. Bourou, T. Boyer, M. Gheisari, K. Daupin, V. Dubreuil, A. De Thonel, V. Mezger, A. Genovesio, "PhenDiff: Revealing Subtle Phenotypes with Diffusion Models in Real Images" in *Medical Image Computing and Computer Assisted Intervention –*

*MICCAI 2024*, M. G. Linguraru, Q. Dou, A. Feragen, S. Giannarou, B. Glocker, K. Lekadir, J. A. Schnabel, Eds. (Springer Nature Switzerland, Cham, 2024), pp. 358–367.

12. C. Gayrard, N. Borghi, FRET-based Molecular Tension Microscopy. *Methods* **94**, 33–42 (2016).

13. R. Rondon, T. Hezez, J. Richard Albert, S. Hayashi, B. Drayton-Libotte, G. G. Curto, F. Auradé, E. Balloul, C. Dugast-Darzacq, F. Relaix, P. Gilardi-Hebenstreit, V. Ribes, Dual transcriptional activities of PAX3 and PAX7 spatially encode spinal cell fates through distinct gene networks. *PLOS Biol.* **23**, e3003448 (2025).

14. M. I. Love, W. Huber, S. Anders, Moderated estimation of fold change and dispersion for RNA-seq data with DESeq2. *Genome Biol.* **15**, 550 (2014).

**Data S1. Sample sizes, statistical tests, and exact P values for all graphs presented in the figures.**

**Data S2. Bulk RNA-seq dataset from day 7 WT, *PAX3*$^{+/-}$, and *PAX3*$^{-/-}$ spinal organoids and gene sets for enrichment analyses.**

**Sheet 1:** Bulk RNA-seq dataset for three independent WT, *PAX3*$^{+/-}$, and *PAX3*$^{-/-}$ clones (clone 1 to clone 3). Column A: Entrez/NCBI gene ID. Column B: Gene symbol. Columns C-K: Fragments per kilobase of transcript per million mapped reads (FPKM). Columns L-T: Transcripts per million (TPM). Columns U-AC: Raw read counts. Columns AD-AL: Normalised read counts. Columns AM-AO: Mean FPKM values across the three biological replicates for each genotype. Columns AP-AX: Log2 fold changes, adjusted p-values (padj), and differentially expressed gene (DEG) status from all pairwise differential expression analyses performed using DESeq2 between the indicated genotypes.
**Sheet 2:** List of genes differentially expressed between the three genotypes, and the cluster to which they belong (k-means, 4 clusters).
**Sheet 3:** Gene sets used for the hypergeometric enrichment analyses shown in Fig. 5B, including gene sets related to dorsoventral (DV) patterning, Wnt signalling, Notch signalling, SHH signalling, glucose metabolism, and cytoskeleton remodelling.
**Sheet 4:** Gene sets used for the ssGSEA analyses shown in Fig. S8D.

**Data S3. Untargeted metabolomics dataset of WT and *PAX3*$^{-/-}$ spinal organoids under the indicated metabolic perturbations (non-treated, mannose supplementation, and glucose deprivation).**
Columns A-R: Metabolite annotation and identification information, including experimental m/z, retention time, annotation confidence, in-house and public database annotations, metabolite identifiers, chemical descriptors, metabolite class, and associated metabolic pathways. Columns S-U: Quality control metrics (pool CV, blank-over-sample ratio, and pool dilution correlation). Columns V-BK: Statistical analyses for each of the seven pairwise comparisons, including Wilcoxon p-values, fold changes (ratio), Benjamini-Hochberg-adjusted p-values, variable importance in projection (VIP) scores, and PCA loadings. Column BL: Chromatographic column used for metabolite detection. Columns BM-EE: Intensity values for each biological sample (six groups of ten replicates) and for the quality control (QC) injections.
